# Substrate-dependent cluster density dynamics in bacterial phosphotransferase system permeases

**DOI:** 10.1101/349514

**Authors:** Gustavo Benevides Martins, Giacomo Giacomelli, Marc Bramkamp

## Abstract

Bacteria take up carbohydrates by membrane-integral sugar specific phosphoenolpyruvate-dependent carbohydrate:phosphotransferase systems (PTS). Although PTS is at the heart of bacterial carbon uptake and centrally involved in regulation of carbon metabolism, little is known about localization and putative oligomerization of the permease subunits (EII) of PTS. Here, we analyzed localization of the fructose specific PtsF and the glucose specific PtsG transporters from *C. glutamicum* using widefield and single molecule localization microscopy. PtsG and PtsF form membrane embedded clusters that localize in a punctate pattern within the cell membrane. The size, number and fluorescence of the observed clusters changes upon presence or absence of the transported substrate. In presence of the transport substrate clusters significantly increased in size. Photo-activated localization microscopy (PALM) data revealed that, in presence of different carbon sources, the number of EII protein events per cluster remain the same, however the density of PTS molecules within a cluster reduces. Our work reveals a simple mechanism for efficient membrane occupancy regulation. Clusters of PTS EII transporters are densely packed in absence of a suitable substrate. In presence of a transport substrate the EII proteins in individual clusters occupy larger membrane areas, thereby decreasing protein density in individual clusters. This mechanism allows for efficient use of the limited membrane space under varying growth conditions without need of protein removal and re-synthesis.

**Importance:** The carbohydrate transport system PTS is centrally involved in the regulation of sugar metabolism. Although much is known about the regulatory interaction, the genetic control and the structure/function relationship of the individual PTS components, we know almost nothing about the spatio-temporal organization of the PTS proteins within the cell. We find dynamic clustering of PTS permeases in *Corynebacterium glutamicum.* Using single molecule resolution photo-activated localization microscopy we could show that PTS EII protein cluster are dynamically changing protein density upon substrate availability. Our findings imply a novel strategy of regulating limited membrane space efficiently. Furthermore, these data will provide important insights in modelling carbohydrate fluxes in cells, since current models assume a homogeneous distribution of PTS permeases within the membrane.

## Introduction

In heterotrophic bacteria, uptake of suitable carbohydrates is an essential task for the cells in their quest for food and, hence, subject to meticulous regulation. In growth media with several carbon sources, many bacteria including enteric bacteria like *Escherichia coli*, use preferred sugars such as glucose first. This leads to the well-known diauxic growth behavior (1, 2). Regulation of sequential carbohydrate usage and transport of carbohydrates is often governed by an enzyme complex termed phosphoenolpyruvate-dependent carbohydrate:phosphotransferase systems (PTS) (3–8). Interestingly, some bacteria are known to co-ferment different carbohydrates with *Corynebacterim glutamicum* being a prominent example (9, 10). *C. glutamicum* is a facultative anaerobic, chemoheterotroph that is used in the large scale industrial production of amino acids (11–13). In industrial applications, the main feedstock used in most of the established fermentation processes are molasses and starch hydrolysates, which contain a broad spectrum of simple carbohydrates, but mostly glucose, fructose and sucrose. These sugars are taken up and phosphorylated during transport into the cell via PTS. This means that worldwide amino acid production using this organism depends heavily on the PTS transport activities (14). The PTS consists of two common energy-coupling cytoplasmic proteins, enzyme I (EI) and histidine protein (HPr), which are encoded by *ptsI* (cg2117) and *ptsH* (cg2121), respectively, and a series of sugar-specific enzyme II (EII) complexes located in the membrane. The EII complexes are typically divided into three protein domains, EIIA, EIIB and EIIC, whose organization differs between bacteria, ranging from organisms in which the three EII domains are fused in a single protein to a variety of differently fused and unfused domains (3, 7). In *E. coli*, the glucose PTS subunit EIIA is detached, while in *B. subtilis* and *C. glutamicum*, all three subunits are fused together. In enteric bacteria EIIA (also termed EIIA^crr^, since the coding gene is abbreviated *crr*), is at the heart of a complex regulatory system that governs the quest for food (4–6). EIIA^crr^ interacts with several transport proteins, such as the lactose permease LacY, thereby inhibiting their activity. This leads to the inducer exclusion effect (15). The accumulation of phosphorylated EIIA^crr^, a sign of starvation, activates the adenylate cyclase, leading to cyclic AMP (cAMP) production. The second messenger cAMP acts as coactivator for the Crp protein cumulating in permanent catabolite repression in enteric bacteria. The passage of specific sugars through the membrane is catalyzed by the transmembrane EIIC part of the system (16). Structural analysis of EIIC subunits revealed that they likely work as dimers. They contain a substrate recognition and a transport part (17). The PTS couple transport to the conversion of carbohydrates into their respective phosphoesters using the energy of phosphoryl group translocation. The phosphoryl group from phosphoenolpyruvate (PEP) is transferred to EI, HPr, EIIA, EIIB and finally to the substrate as it is transported across the membrane. In *C. glutamicum* four PTS transport system have been described. The organization of the *C. glutamicum* PTS for fructose, glucose and sucrose are represented in figure 1. *ptsG* (cg1537) encodes the glucose-specific membrane integral EIIBCA transporter, while *ptsS* (cg2925) encodes the sucrose-specific EIIBCA, and *ptsF* (cg2120) the fructose-specific EIIABC. A fourth PTS belongs to the L-ascorbate-type family and was characterized by R. Hvorup, et al. (18)(not depicted in the graphic).

**Figure 1.**
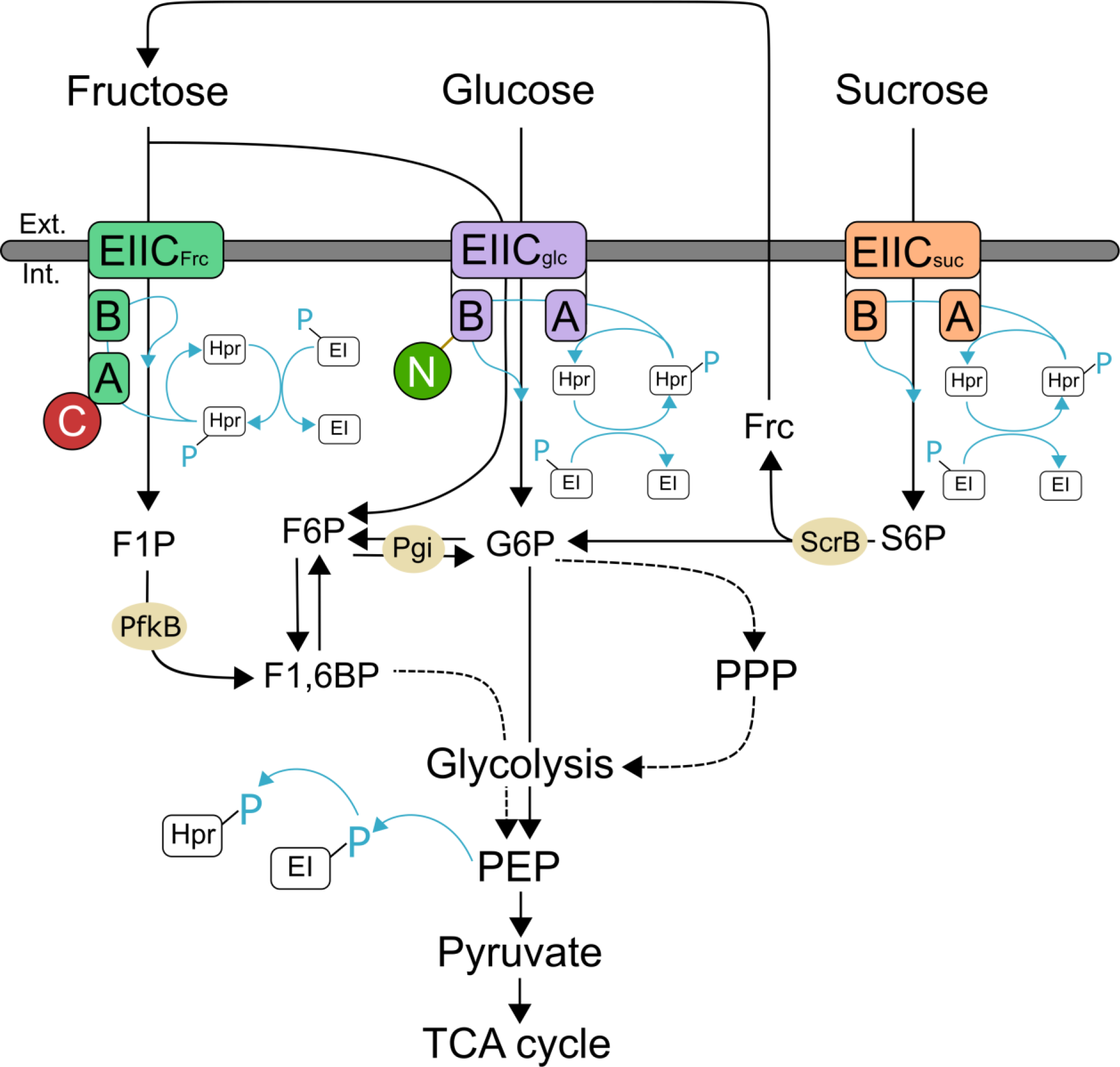
Schematic view of the sugar uptake mediated by the phosphoenolpyruvate-dependent phosphotransferase systems (PTS) in *Corynebacterium glutamicum.* Blue lines indicate phosphoryl group (PO_3_^2-^) transfer. Fluorescent proteins added via allelic replacement represented as C for mCherry in subunit EIIA of PtsF, and N for mNeonGreen-Linker in subunit EIIB of PtsG. F1P *fructose-1-phosphate*, F6P *fructose-6-phosphate*, F1, 6BP *fructose-1,6-*870*biphosphate*, G6P *glucose-6-phosphate*, Frc*fructose*, S6P *sucrose-6-phosphate*, PPP *Pentose phosphate pathway*, PEP*phosphoenolpyruvate*, scrB *beta-fructofuranosidase* (putative S6P hydrolase), pgi *glucose-6-phosphate isomerase*, 873PfkB *1-phosphofructokinase*.

EII expression is known to be induced by the presence of its transported sugar (19), and regulation of PTS gene expression is mainly controlled at the stage of transcription initiation or at transcription elongation in *C. glutamicum* (19). The mechanism for control of PTS gene expression differs for the respective PTS genes. The transcriptional regulator SugR, encoded by *sugR* (cg2115) is a deoxyribonucleoside repressor (DeoR)-type regulator and is located in the fructose-PTS operon. It has been shown to repress not only *ptsF*, *ptsG* and *ptsS* expression in the absence of the transported sugar, but also the general PTS genes *ptsI* and *ptsH* in *C. glutamicum* (20–22). Post-transcriptional regulation of PtsG in *E. coli* is governed by SgrS, a small Hfq-binding RNA, that is induced by phosphor-sugar accumulation in the cytosol. It forms a ribonucleo-protein complex with Hfq and RNaseE, resulting in translational repression and degradation of *ptsG* mRNA (23, 24).

The catabolite repressor/activator FruR (also known as Cra), encoded by *fruR* down regulates the fructose-PTS operon in *E. coli, Salmonella typhimurium* and *Pseudomonas putida* when the sugar is not available (22, 25, 26). In *P. putida*, FruB acts by binding as a dimer to a single palindromic sequence within the *cra/fruB* intergenic region (which is different to those of *E. coli* and S. *typhimurium*), and that only fructose-1-phosphate can lift such down regulation (26). In *S. pneumonia*, exogenous auto-inducer 2 (AI-2) induces *fruA* and *fruB*, which encode PtsF and 1-phosphofructokinase in that organism (27). Expression of enzyme I and HPr increases in the presence of different PTS sugars. In *Streptomyces coelicolor* and *Lactobacillus casei*, glucose is reported to be the most effective inducing sugar of *ptsI* and *ptsH* expression (28, 29). In *C. glutamicum*, the role of FruR (cg2118) is distinct from the one observed in other bacteria. It represses the transcription of the *fruR-pfkB1-ptsF* operon, as well as *ptsI* and *ptsH* in presence of fructose, decreasing the induction effect of fructose and adjusting the expression level of *pts* genes to prevent overflow of PTS sugars (30)

It has been proposed that the PTS can be described as a central “cellular functioning unit” [CFU], regulating the cells quest for food (4–6). However, to fully describe CFUs, precise knowledge about spatio-temporal localization of its components is required. There is surprisingly little knowledge on subcellular localization of PTS components. Early immune-gold labelling with the *E. coli* EII^mtl^ showed membrane localization that can be interpreted as patchy with no preferred subcellular enrichment (31). A more recent study showed that the *E. coli* BglF (the β-glucoside specific EII permease) is located throughout the plasma membrane (32). However, deconvolved images and plasmid born expression do not allow for unambiguous differentiation between a uniform and a patchy localization. The general components of the *E. coli* PTS, EI and HPr in contrast localize to the cell poles (32, 33). HPr localization changes in presence of the transport substrate from polar to dispersed throughout the cytoplasm.

Here, we describe the localization of two specific EII components in *C. glutamicum.* We have constructed translational fusions of *ptsG* and *ptsF* encoding PtsG (EII^glc^) and PtsF (EII^frc^) with different fluorescent proteins. Constructs were designed as allelic replacements, ensuring native genetic control and all constructs were shown to be fully functional. Widefield fluorescence microscopy revealed that EII complexes localize as dynamic clusters in the cell membrane. PtsG and PtsF EII components exclude each other within the membrane compartment, but PtsF co-localize with components of the respiratory chain, ruling out a specific membrane domain for carbohydrate transport only. Importantly, we observed an increase in PTS EII cluster size when the specific sugar substrate was present. This increase in cluster sized coincided with a decrease in cluster number. Using photo-activated single molecule fluorescence microscopy (PALM) we were able to quantitatively address PTS dynamics. PALM data clearly show that PTS EII cluster are covering a larger membrane area when their transport substrate is presence. Importantly, under these conditions the complexes do not contain more EII molecules, but rather reduce protein density within clusters. Thus, actively transporting PTS permeases are spreading apart and non-transporting complexes are densely packed. This dynamic arrangement of the PTS offers a simple mechanism for efficient membrane occupancy under varying nutrient conditions.

## Results

### Construction of functional PTS fusion proteins

To investigate subcellular localization and membrane occupancy of PTS EII permeases, we constructed *C. glutamicum* strains with fluorescent fusions to PtsF and PtsG, resulting in strains CGM001 and CGM002 expressing mCherry-PtsF and mNeonGreen-PtsG, respectively (Table S2). We have not included the sucrose specific PTS since sucrose is a disaccharide composed of fructose and glucose and we wanted to first test for the specific influence of glucose and fructose. *C. glutamicum* is importing sucrose via the sucrose specific PTS. The phosphorylated sucrose is cleaved intracellularly and the resulting fructose molecule is exported and taken up by the fructose specific PtsF. Hence, a clean separation between sucrose and fructose effects is not easily possible. To check for potential colocalization of the glucose and fructose specific EII permeases a double labelled strain was constructed. To this end, CGM001 was used as a background for incorporation of *mNeonGreen-linker-ptsG* via allelic replacement, resulting in the double-labelled strain *ptsF::mCherry-ptsF, ptsG:.mNeonGreen-linker-ptsG* (CGM003) (Fig. 2A). In order to gain quantitative single molecule resolution data of PTS localization and clustering behavior, strains containing PtsF (CGM004) and PtsG (CGM005) tagged with photoactivatable mCherry (PAmCherry) were constructed for PALM microscopy. To avoid overexpression and artificial expression heterogeneity, the fusion constructs were all inserted as allelic replacement in the genome of wild type *C. glutamicum* (RES 167) cells. To test functionality of the constructed fusion proteins, growth experiments and sugar consumption assays via HPLC were performed. Strains CGM001, expressing mCherry-PtsF (Fig. 2B), CGM002 expressing mNeonGreen-PtsG (Fig. 2C) and the double labelled strain CGM003 displayed wild type like growth behavior (Figs. 2D and 2E). The growth rates of wild type did not differ from the constructed strains and ranged from 0.28 to 0.29 h^-^1 (Tab. 1), showing that the tagged PTS systems were functioning wild type-like. We also measured sugar consumption rates in CGXII supplemented with 2% fructose or 2% glucose. The specific sugar consumption rates qs were highly similar for wild type and mutant strains (Tab. 1). The measured qs values for the wild type of around 0.19 g g^−1^ h^−1^ for glucose and 0.13 g g^−1^ h^−1^ for fructose are very well within the expected range for the observed growth rate (34). The strain carrying the fluorescently labelled PtsG had consumption rates of 0.18 g g^−1^ h^−1^ for glucose and the strain with the PtsF fusion construct displayed consumption rates of 0.14 g g^−1^ h^−1^ for fructose. In summary we conclude that the constructed fusions are fully functional. It should be noted here, that in a process to gain functional translational fusions, several constructs were made that were all not further analyzed when they turned out to be non-functional.

**Table 1.** Growth rates and glucose consumption rates. Growth rate [h^−1^] and substrate uptake [g g-1 h-1] of RES 167 WT and strains CGM001, CGM002, CGM003 and CGM005 in CGXII supplemented with glucose and fructose.

|  | Glucose |  | Fructose |  |
| --- | --- | --- | --- | --- |
| | growth rate [$\text{h}^{-1}$] | uptake (qs) [$\text{g g}^{-1} \text{h}^{-1}$] | growth rate [$\text{h}^{-1}$] | uptake (qs) [$\text{g g}^{-1} \text{h}^{-1}$] |
| **WT** | 0.285 | $0.23 \pm 0.003$ | 0.289 | $0.14 \pm 0.005$ |
| **CGM001** | - | - | 0.286 | $0.17 \pm 0.004$ |
| **CGM002** | 0.280 | $0.23 \pm 0.001$ | - | - |
| **CGM003** | 0.290 | $0.23 \pm 0.001$ | 0.280 | $0.17 \pm 0.004$ |
| **CGM005** | 0.285 | $0.23 \pm 0.001$ | - | - |

**Figure 2.**
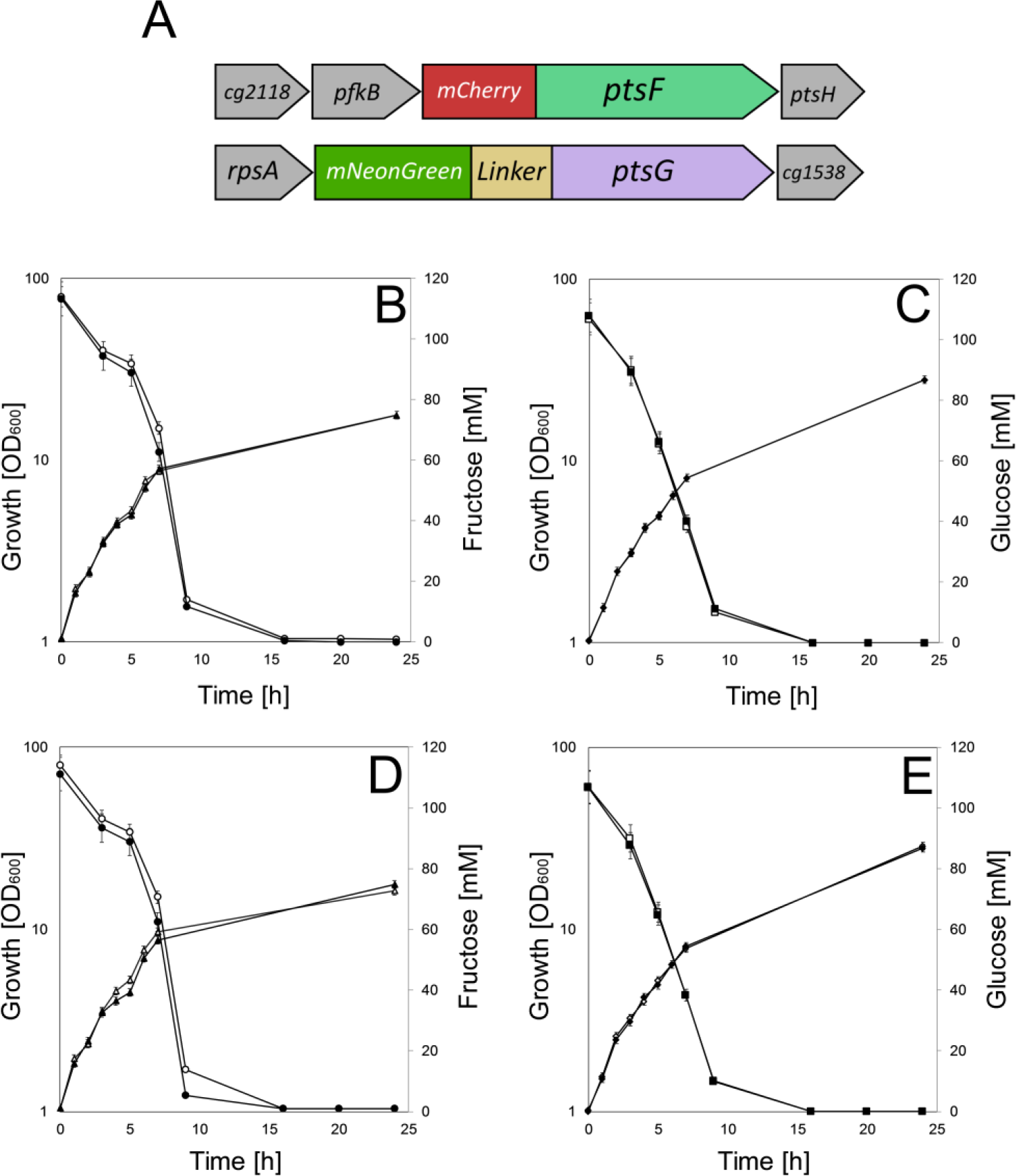
Fluorescent fusions of PtsF and PtsG are fully functional. (A) Genetic arrangement of fluorescent fusions constructed via allelic replacement of *ptsF::mCherry-ptsF* and *ptsG::mNeonGreen-linker-ptsG.* Growth and sugar consumption of *C. glutamicum* strains CGM001, CGM002 and CGM003 (filled symbols) versus wild type RES 167 (open symbols) on CGXII containing (B, D) 100 mM fructose and (C, E) 100 mM glucose. Fructose consumption (circles), glucose consumption (squares), growth on fructose (triangles), and growth on glucose (diamonds) are indicated. Each point represents biological triplicates and standard deviation is indicated.

Protein localization studies using translational fusions can be hampered by protein degradation and subsequent imaging of free fluorophore. Therefore, we investigated protein degradation by western blotting with anti-mCherry for mCherry and PAmCherry fusions, and in-gel fluorescence for mNeonGreen fusions. For each strain, it was possible to identify one the band corresponding to their full length fusion protein, or oligomers, revealing that no major degradation was present (Fig. S1). We therefore conclude that localization studies with these strains should reveal the native localization of the full length PTS EII permeases.

### *C. glutamicum* EII^frc^ EII^glc^ spatial dynamics upon presence or absence of the transported sugars

We first wanted to investigate whether PTS EII proteins distribute uniformly or in clusters along the cytoplasmic membrane under aerobic conditions in presence of the transported sugars. To this end, fluorescence microscopy was performed on *C. glutamicum* strains expressing mNeonGreen-PtsG and mCherry-PtsF grown in CGXII supplemented with the transported sugars as sole carbon sources. Analysis of fluorescence images showed that both proteins form membrane embedded clusters that localize punctually within the cell membrane with no preferred positions (Fig. 3A-B). EII^fru^ and EII^glc^ foci of varying intensity were observed, suggesting complexes with different amounts of proteins. Most cells contained few bright and intense foci that were randomly distributed. However, bright foci seemed to be present with higher frequency close to cell poles (Fig. 3A-B).

**Figure 3.**
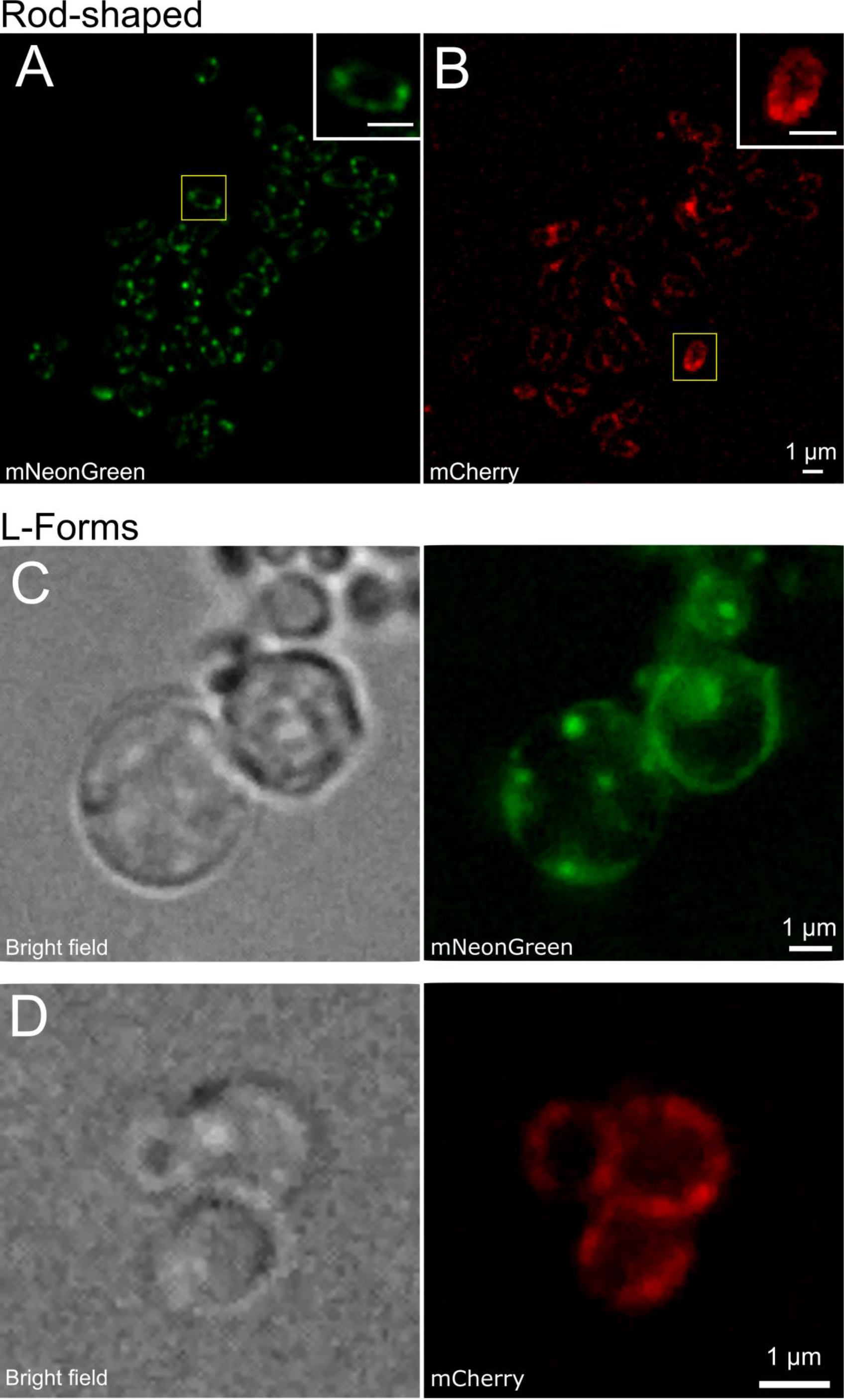
PtsF and PtsG localize as clusters distributed along cytoplasmic membrane. Fluorescence microscopy images of *C. glutamicum* expressing mCherry-PtsF and mNeonGreen-PtsG under different growth conditions: Rod-shaped cells grown in CGXII with (A) glucose and (B) fructose as sole carbon sources. L-Form cells grown in MXM/CGXII with (C) glucose and (D) fructose as sole carbon sources in presence of 200 μg/ml of DCS.

We next wanted to test whether cell shape and the surface/volume ratio might have an influence on the PTS behavior. Therefore, we analyzed PTS localization in L-form bacteria, generated out of the corresponding strains. Cells expressing mNeonGreen-PtsG and mCherry-PtsF were grown in MXM/CGXII medium supplemented with the transported sugars in presence of D-cycloserine (DCS). DCS is a cyclic analogue of D-alanine, acting against alanine racemase (Alr) and D-alanine:D-alanine ligase (Ddl), two crucial enzymes in the cytosolic stages of peptidoglycan synthesis, bypassing the need to block cell wall synthesis genetically. As DCS was previously used in *Mycobacterium tuberculosis* (35), which shares the characteristic cell wall common to all *Corynebacterineae*, it was chosen for L-form formation with *C. glutamicum*. L-form bacteria require an osmotically stabilized medium. Usually, the osmoprotective environment would be achieved by adding sucrose to the media, but since this sugar is taken up by the sucrose specific PtsS, we complemented the L-form medium with xylose, a carbohydrate that cannot be efficiently metabolized by *C. glutamicum*. Fluorescence microscopy with L-forms revealed the same PtsF and PtsG clustering that was observed in rod-shaped cells (Fig. 3C-D). We conclude from this, that cluster formation of PTS EII permeases may be an intrinsic property of these enzymes and not depend on cell shape or surface/volume ratios.

We next wanted to investigate whether presence or absence of the specific PTS carbohydrates influence the PTS EII complexes localization and expression. To this end, the strains containing fluorescent reporters linked to the EII complexes were grown in CGXII minimal medium with 2% glucose, fructose or acetate as sole carbon sources to early exponential phase, and fluorescent microscopy was subsequently performed. Importantly, both proteins are expressed in the absence of their specific transported sugars (Fig. 4 A-C) and there were no differences among early, mid and late log phases (data not shown). Observed PtsG/F foci did not show evident colocalization (Fig. 4D), suggesting that phosphotransferase systems localize independently within the membrane. This is an important observation, because the general PTS components EI and HPr are required for both EII complexes. As a control we analyzed localization of the EII^fru^ complexes with a protein from the respiratory chain. Here, we have chosen the succinate dehydrogenase subunit A (SdhA) in combination with PTS EII. We did observe a large degree of co-localization between PtsF and SdhA (Fig. 4E). These data rule out that in *C. glutamicum* components of the respiratory chain and the PTS would occupy specific and different membrane domains.

**Figure 4.**
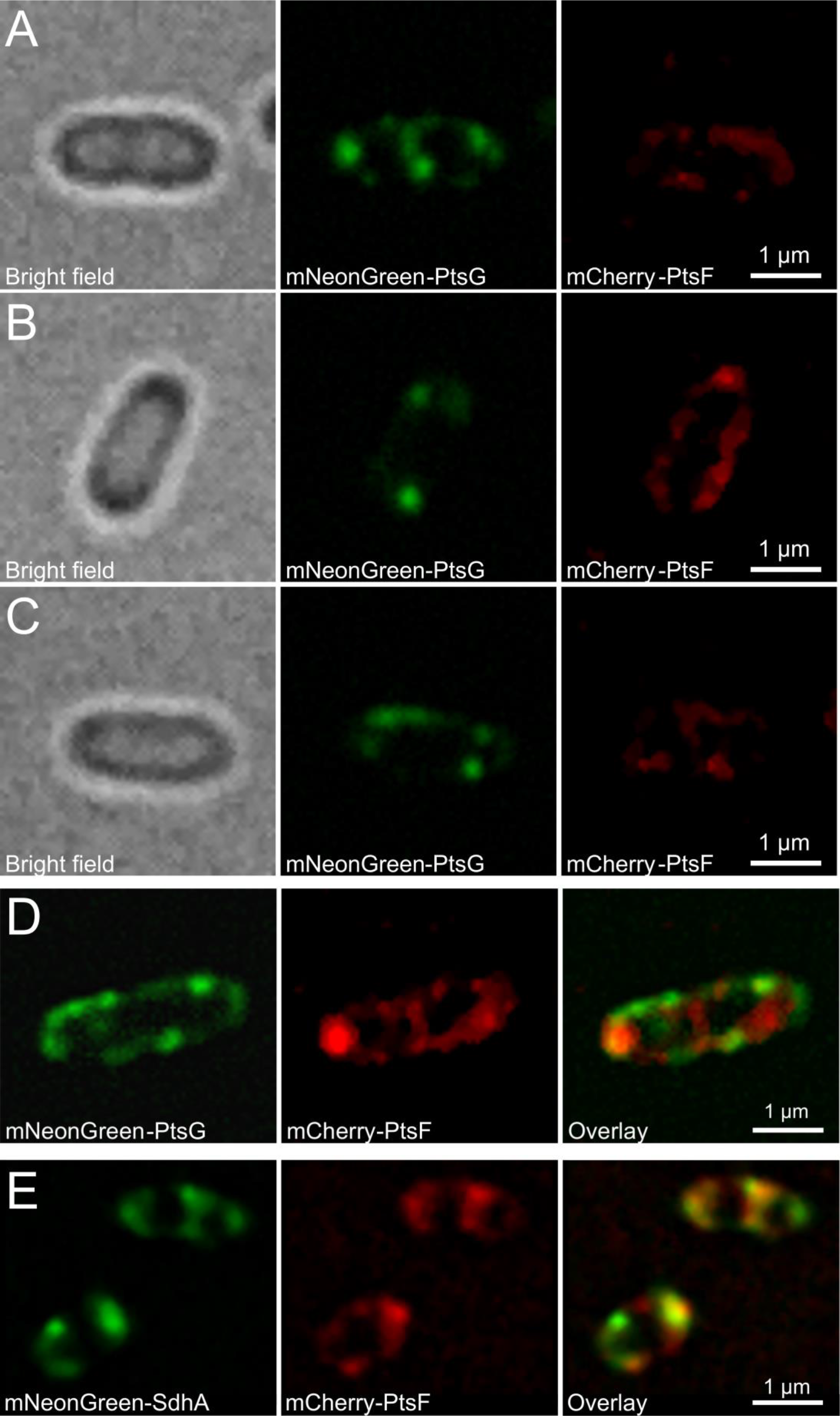
PtsF and PtsG clustering changes upon presence or absence of transported substrate. Fluorescence microscopy images of *C. glutamicum* expressing mCherry-PtsF and mNeonGreen-PtsG under different growth conditions. Cells grown in CGXII with 2% (A) glucose, (B) fructose or (C) acetate as sole carbon sources. (D) mNeonGreen-PtsG and mCherry-PtsF in CGXII with 2% glucose and fructose. (E) mNeonGreen-SdhA and mCherry-PtsF in CGXII with and fructose.

After the observation that clustering and expression of PTS EII occurs even in absence of the transported substrate, we next wanted to determine the influence of different carbon sources on PTS foci number per cell, foci area, fluorescence, and how much of the membrane space is covered by PTS. Analysis of fluorescent images of *ptsF::mCherry-ptsF, ptsG::mNeonGreen-Linker-ptsG* both in rod shaped cells and L-forms grown in minimal medium under different carbon sources revealed that the number of PtsF/G clusters per cell was significantly decreased in presence of the transported sugar in rod-shaped cells (Fig. 5 and Tab. 2). Statistically, the number of PtsF clusters per cell was equal in acetate and glucose treatments, and differed in rod-shaped and L-form cells grown in fructose. Regarding PtsG foci number per cell in rod-shaped, cells grown in fructose or acetate did not differ statistically, whilst when grown in glucose, the values were significantly lower. L-form cells showed a tendency to have more PTS foci, having higher averages (3.3 per cell for both PtsF/G), being statistically different from all the other treatments. The absence of a cell wall in L-forms results in significantly larger cells (mean cell area values: rod-shaped 3.6 μm^2^, L-forms 8.9 μm^2^), which leads to a drastic reduction in the cell surface/volume (S/V) ratio. Despite being grown in medium containing the same carbon source, the observed increase in number of PtsF/G clusters in L-forms suggests that PTS EII foci pattern can be changed in response to variations in cell area, volume or morphology.

**Table 2.** Number of PTS EII foci per cell in various growth conditions. Mean values of foci number per cell of mCherry-PtsF and mNeonGreen-PtsG. “Fructose”, ‘Glucose” and “Acetate” represent rod-shaped cells in CGXII medium with the respective carbon sources. “L-Forms” represent L-Form cells in MXM/CGXII upplemented with the transported sugar. Significant statistical differences according to multiple comparison tests after Kruskal-Wallis are represented as letters next to ach value.

|  | Fructose | Glucose | Acetate | L-Forms |
| --- | --- | --- | --- | --- |
| **PtsF** | 2.4 (b) | 2.6 (a) | 2.9 (a) | 3.3 (c) |
| **PtsG** | 2.8 (a) | 2.5 (b) | 2.8 (a) | 3.3 (c) |

**Figure 5.**
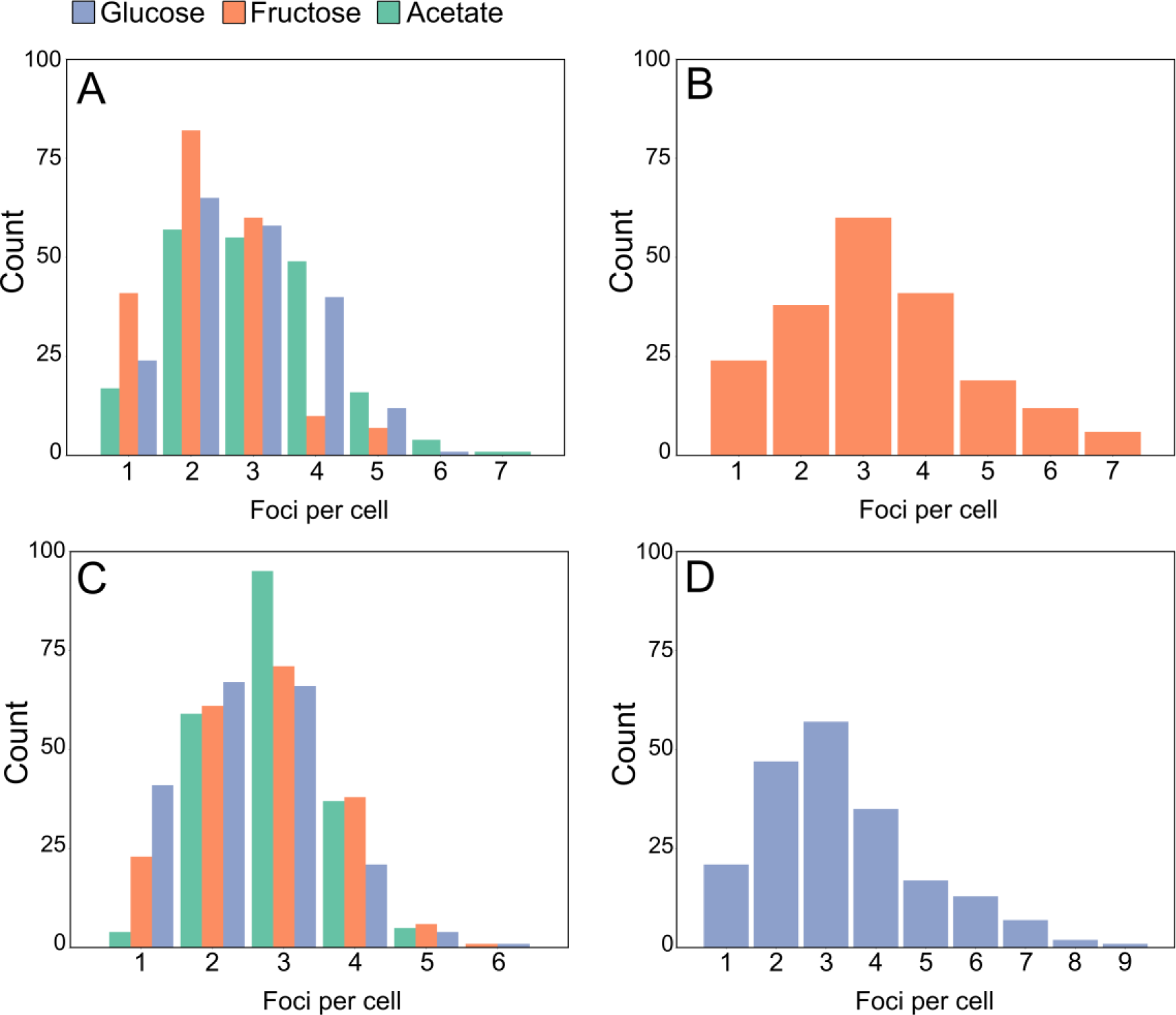
mCherry-PtsF and mNeonGreen-PtsG foci number decreases in presence of transported sugar. Number of foci per cell of (A) mCherry-PtsF in rod shaped cells, (B) mCherry-PtsF in L-form cells. (C) mNeonGreen-PtsG in rod shaped cells, (D) mNeonGreen-PtsG in L-form cells.

Previous studies have revealed that the expression of the *C. glutamicum* PTS is induced when cells are cultured in presence of the transported sugar (19–21, 36). In order to test the influence of different carbon sources on PTS foci, PtsF/G Corrected Total Fluorescence (CTF) was calculated with ImageJ based on the obtained fluorescence microscopy images and corrected for the cell area (Fig. 6A,B). Analysis of CTF of tagged proteins served as a proxy to assess levels of protein expression and concentration, and both PtsF and PtsG were similar regarding the fluorescence readings in this work. Although all the treatments were statistically different from each other, CTF of both proteins increased in presence of the transported substrate and decreased in its absence. This indicates that indeed, PtsF and PtsG expression was induced in presence of the transported substrate. When compared to rod shaped cells grown in the same carbon source, the CTF of L-forms was over 4 times higher (4,5× for PtsF and 4,1× for PtsG), this suggests that the number of PTS EII complexes within the cell membrane increases with the cell area and number of chromosomes, since large L-forms are known to larger numbers of chromosomes (37).

**Figure 6.**
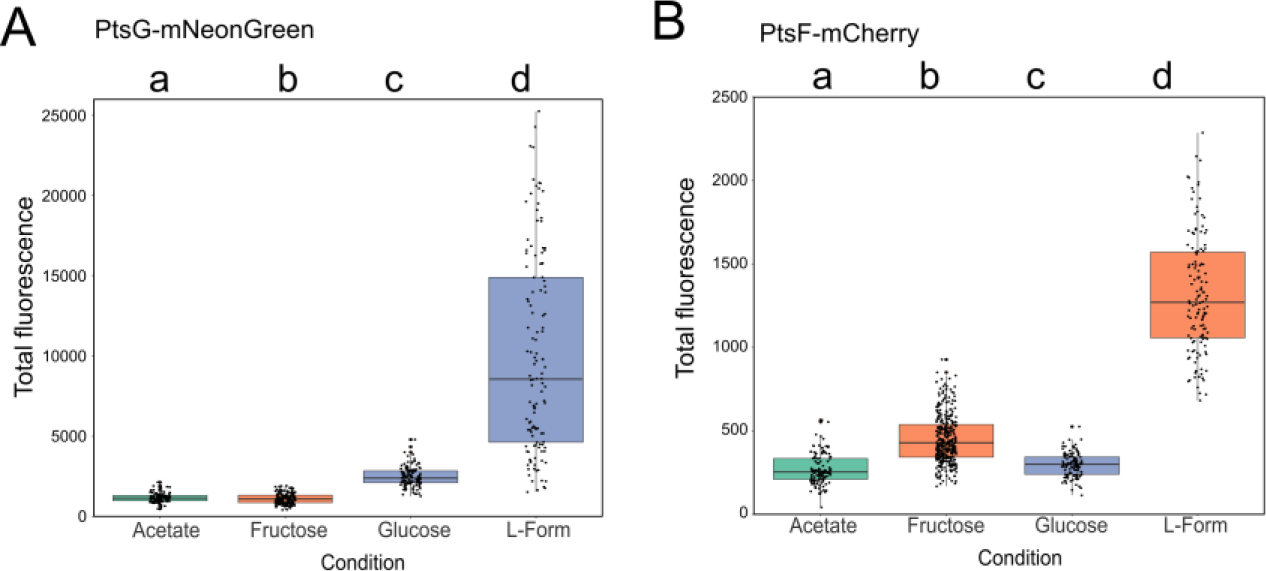
*C. glutamicum* PTS expression increases in presence of transported sugar. Corrected total fluorescence of (A) mNeonGreen-PtsG and (B) mCherry-PtsF. “Fructose”, “Glucose” and “Acetate” represent rod-shaped cells in CGXII medium with the respective carbon sources. “L-form” represents L-form cells in MXM/CGXII supplemented with fructose for PtsF and glucose for PtsG. Significant statistical differences according to multiple comparison tests after Kruskal-Wallis are represented as letters above each graph.

The areas of the PtsF/G foci of cells grown in different carbon sources are summarized in Figure 7A and Table 3. In rod-shaped cells, PtsF foci area values were statistically similar when cells were grown in glucose or acetate, and significantly higher in fructose. Likewise, PtsG foci area in glucose was 3 times higher than in fructose, and 4 times than in acetate. However, unlike PtsF, the PtsG cluster area values obtained for cells grown in fructose were higher than cells grown in acetate. L-Form cells exhibited the highest PtsF/G foci areas compared to every other condition in rod-shaped cells, suggesting that a larger membrane area create more space to be occupied by transmembrane proteins.

**Table 3.** PtsF and PtsG cover different membrane surface areas. Mean values of foci area [μm^2^] of mCherry-PtsF and mNeonGreen-PtsG. “Fructose”, “Glucose” and “Acetate” represent rod-shaped cells in CGXII medium with different carbon sources. “L-Forms” represent L-Form cells in MXM/CGXII supplemented with the transported 33sugar. Values are in μm^2^. Significant statistical differences according to Multiple comparison tests after Kruskal-Wallis are represented as letters next to each value.

|  | Fructose | Glucose | Acetate | L-Forms |
| --- | --- | --- | --- | --- |
| **PtsF** | 0.35 (b) | 0.18 (a) | 0.24 (a) | 0.71 (c) |
| **PtsG** | 0.22 (a) | 0.70 (b) | 0.17 (c) | 1.00 (d) |

**Figure 7.**
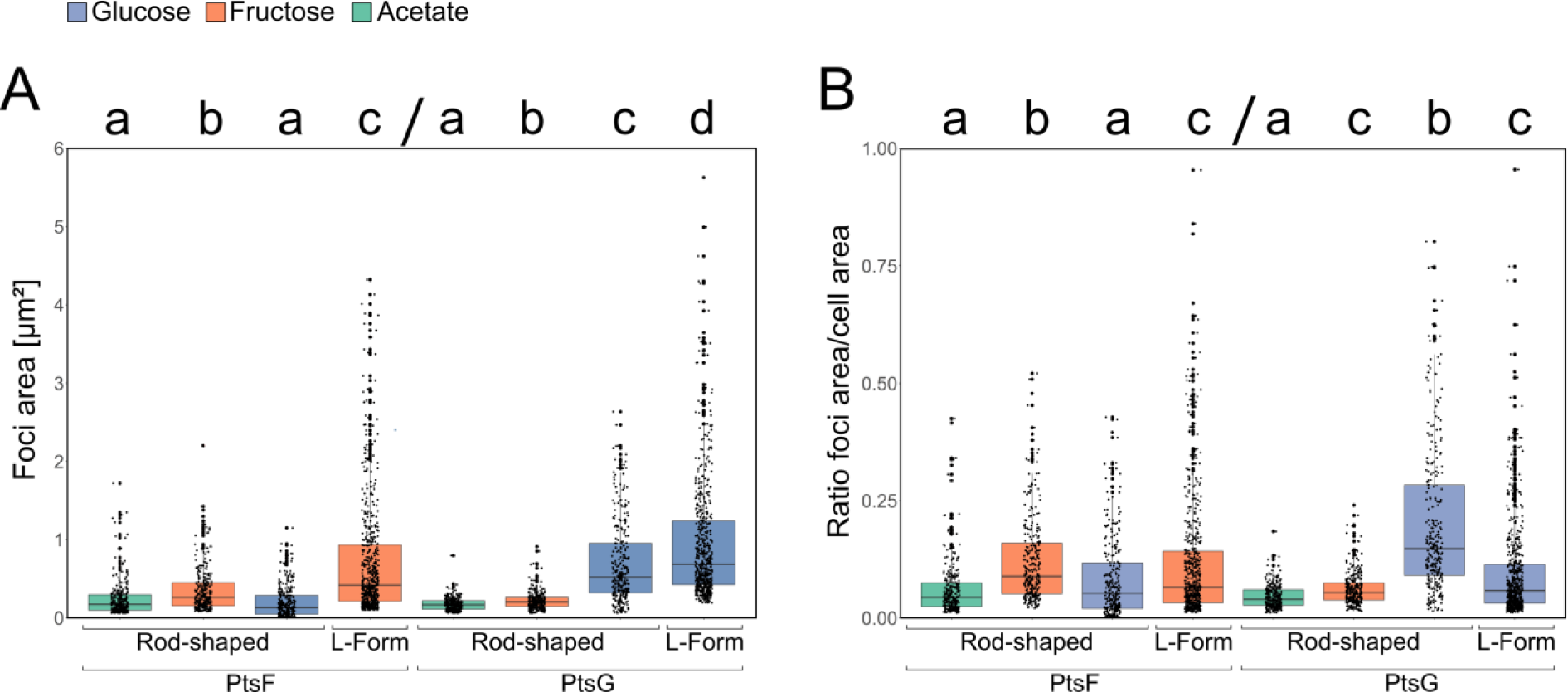
mCherry-PtsF and mNeonGreen-PtsG foci area increases upon presence of transported sugar. (A) mCherry-PtsF and mNeonGreen-PtsG fluorescent foci area in rod-shaped and L-Form cells in CGXII supplemented with different carbon sources (B) Ratio foci area and cell area under different carbon sources and cell shapes. “L-Form” represent L-Form cells in MXM/CGXII supplemented with the transported sugar. Significant statistical differences according to multiple comparison tests after Kruskal-Wallis are represented as letters above each graph. Different letters indicate differences within the same PTS protein.

In order to know how much of the cytoplasmic membrane is occupied by PTS, the ratio foci area/cell area (Fig. 7B) was calculated. Rod-shaped cells had the highest ratios when in presence of the transported sugar, meaning that more membrane space is reallocated for PTS proteins in such conditions. In average, PtsG ratio in presence of glucose was 3 times higher than in fructose, and 4 times than in acetate. Although less dramatic, the increase in PtsF ratio while in presence of fructose was still significant: 1,8 times higher than in glucose, and 1,45 higher than acetate. In general, L-forms exhibited larger PTS clusters, in a higher overall number, and fluorescence. This leads to the logical assumption that a larger cell area increases the area available for protein insertion, possibly resulting in more transmembrane PTS proteins. However, the foci area/cell area ratio of L-Forms was not higher than other rod-shaped cells. In fact, despite PtsF L-form values being higher than in rod-shaped cells grown in glucose or acetate, they were lower than rod-shaped cells in fructose (Table 4). Roughly, the same was observed for PtsG: even though L-forms had a higher foci area/cell area ratio than rod-shaped cells in general, there was no statistical difference between their values and rod-shaped grown in fructose, a condition where PtsG is hardly induced.

**Table 4.** PTS EII complex surface coverage. Foci area/cell area ratio of mCherry-PtsF and mNeonGreen-PtsG. “Fructose”, “Glucose” and “Acetate” represent rodshaped cells in CGXII medium with different carbon sources. “L-Forms” represent L-Form cells in MXM/CGXII supplemented with the transported sugar. Significant statistical differences according to multiple comparison tests after Kruskal-Wallis are represented as letters next to each value.

|  | Fructose | Glucose | Acetate | L-Forms |
| --- | --- | --- | --- | --- |
| **PtsF** | 0.10 (b) | 0.05 (a) | 0.06 (a) | 0.08 (c) |
| **PtsG** | 0.06 (c) | 0.20 (b) | 0.04 (a) | 0.11 (c) |

### Single molecule localization microscopy reveals spatial rearrangement of EII^glc^ in presence of glucose

The data obtained with widefield microscopy suggested that PTS EII cluster rearrange when their specific transport substrate is present. However, epifluorescence microscopy is limited by the diffraction limit, and a series of fundamental information about PTS EII clusters such as density or number of molecules cannot be obtained. Therefore, the next step we took towards a deeper and more quantitative understanding of PTS EII complex dynamics was to use single molecule photo-activated localization microscopy (PALM) data. To this end, the construction of strains tagged with photoactivatable mCherry (PAmCherry) *ptsG::PAmCherry-Linker-ptsG* and *ptsF::PAmCherry-ptsF* was carried out the same way as the previous strains and was also shown to be fully functional as judged by sugar uptake and growth rates (Fig. S2). The observed clustering pattern in epifluorescence was confirmed by PALM data of cells expressing PAmCherry-PtsG (strain CGM005) (Fig. 8 A,B). Also, PAmCherry-PtsF formed membrane embedded clusters, but the low number of events detected per cell made it impossible to obtain significant statistical analysis that would show cluster density changes (Fig. S3). However, we observed a tendency that is in accord with the data obtained for PtsG.

**Figure 8.**
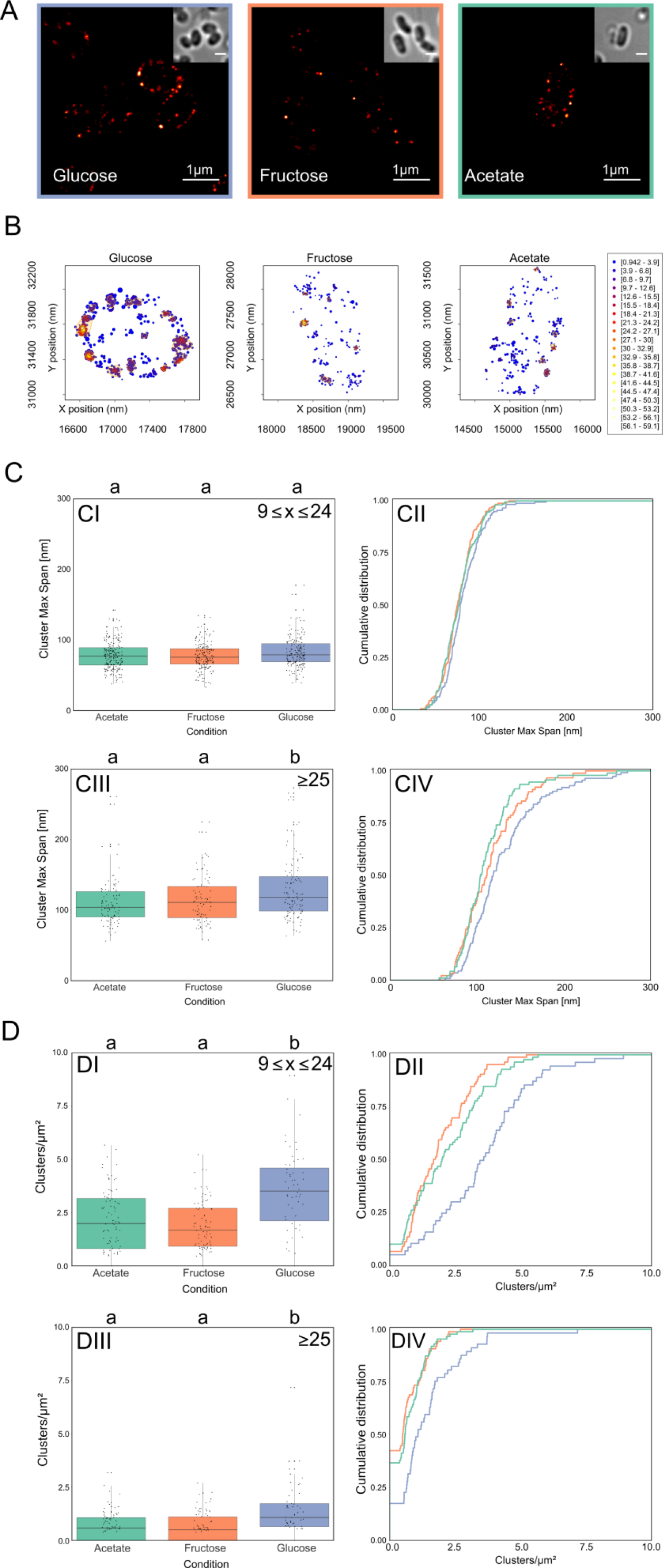
Single molecule localization microscopy reveals dynamic PAmCherry-PtsG clustering. (A) Super resolution images of *C. glutamicum* strain CGM005 expressing PAmCherry-PtsG under different growth conditions. Cells were grown in CGXII with indicated carbon source. Insets show transmitted light images to see cell outlines. Scale bar is 1 μm. (B) Plot of detected single molecule localization events localization and clusters identification for a representative cell for each tested condition. Localizations are color coded according to the local event density while the plotted radius is indicating localization precision. The identified clusters are highlighted by orange contours. (C) PtsG clusters maximum span increase in presence of glucose, (D) PtsG clusters are more abundant in presence of glucose. Each parameter is analyzed via (CI, CIII, DI, DIII) boxplot and (CII, CIV, DII, DIV) cumulative distribution function. Clusters composed of (CI, CII, DI, DII) 9 ≤ x ≤ 24 events, and (CIII, CIV, DIII, DIV) >25 events were analyzed. Significant statistical differences according to multiple comparison tests after Kruskal-Wallis are represented as letters above each graph. Different letters indicate statistical differences within the same PTS protein.

For analysis of PALM data we first needed to precisely define what a valid cluster is. Clusters are defined as regions of high density separated by regions of lower density, and the distribution our data suggests is that there are three populations separated by the amount of events composing each cluster: (A) X>10, (B) 10≤X≤24, and (C) X≥25 events (Fig. S4). The first population could in theory still be composed of PTS complexes close to each other by mere coincidence. Although this randomness effect is never fully absent, it decreases with increasing cluster size. Therefore, clusters composed of ≥10 and ≥25 events were analyzed regarding their maximum span, density, number of events per μm^2^, and number of clusters per μm^2^. The stretched exponential distribution of PTS cluster size is reminiscent to that observed with chemotaxis receptors (38) and therefore suggests a stochastic self-assembly process. We detected around 177.65 (sd = 92.48) PTS EII^glc^ events per μm^2^ cell area in glucose, 111.22 (sd = 49.51) events per μm_2_ in fructose, and 114.69 (sd = 60.28) events per μm^2^ in acetate.

The increase in foci area observed in epifluorescence brought up the question whether the cluster area increases due to an increase in number of PTS complexes present in each cluster, or to a rearrangement of the same number of EII permeases per cluster. At single molecule resolution, foci area can be estimated by the cluster maximum span (Fig. 8C), which is the maximum distance between two events belonging to the same cluster. PAmCherry-PtsG clusters from populations B and C exhibited significantly higher max span under the presence of glucose when compared to fructose or acetate, which exhibited no statistical difference among each other, corroborating our findings with epifluorescence, where in presence of the transported sugars, PtsG/F covered a larger membrane area. The analysis of these same populations of clusters revealed that cells in presence of glucose exhibited significantly higher number of PtsG clusters per μm^2^ when compared to fructose and acetate, which exhibited no differences among each other (Fig. 8D). CTF readings of epifluorescence images showed increased values when cells expressing mCherry-PtsF and mNeonGreen-PtsG were grown in presence of the transported sugars, suggesting increased expression of PTS complexes under these conditions. This data correlates with our PALM data for PamCherry-PtsG (Fig. 9A). In presence of glucose, a significantly higher number of events per μm^2^ was observed, meaning that the number of EII^glc^ in the cytoplasmic membrane increases with the addition of glucose in the medium, again confirming induction of *ptsG* in presence of glucose as expected. Our epifluorescence data suggested that in presence of the transported sugar, PtsF/G assemble in larger complexes, while the overall number of clusters per cell decreases. However, our PALM analysis of cells expressing PAmCherry-PtsG in glucose showed an increase in the number of clusters and events per μm^2^. Single molecule detection allowed visualization of clusters with fewer proteins, thereby explaining the apparent discrepancy to the widefield microscopy data. While in widefield the number of bright, visible foci decreased (while their area increased) when the transport substrate was present, in PALM we observed more clusters with lower protein numbers that likely account from substrate induced gene expression.

**Figure 9.**
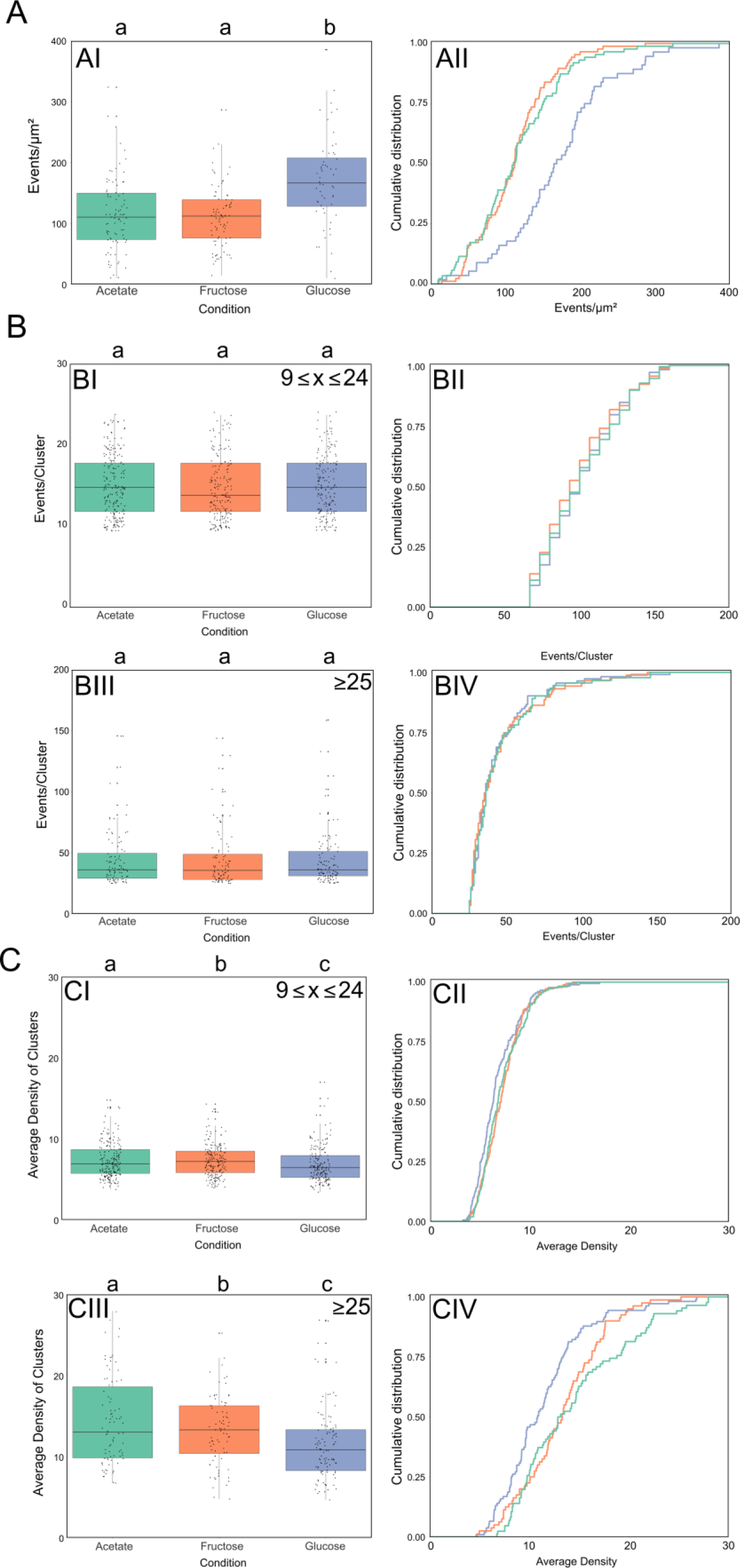
PTS cluster decrease protein density in presence of the transport substrate. Super resolution PALM data of CGM005 strain grown in CGXII supplemented with 2% of the indicated carbon sources (A) PtsG number of events per μm^2^ increases in presence of glucose, (B) PtsG clusters are composed by the same amount of molecules regardless of the carbon source. Plots represent events per cluster in clusters composed of >25 molecules. (C) PtsG clusters have lower average density in presence of glucose. Each parameter is analyzed via (AI, BI, BIII, CI, CIII) boxplot and (AII, BII, BIV, CII, CIV) cumulative distribution function. Clusters composed of (BI, BII, CI, CII) 9 ≤ x ≤ 24 events, and (BIII, BIV, CIII, CIV) >25 events were analyzed. Significant statistical differences according to multiple comparison tests after Kruskal-Wallis are represented as letters above each graph. Different letters indicate statistical differences within the same PTS protein.

Although the CTF of cells increases in presence of glucose, PALM data showed that the number of PAmCherry-PtsG events per cluster remain the same independently of the carbon source, suggesting that the carbon source does not affect the number of PTS proteins present in each cluster (Fig. 9B). This is an important finding since larger cluster size could have been the trivial consequence of more protein in the cell. Since we have found that the number of PTS EII molecules in a cluster remained similar in presence or absence of the transported substrate, we wanted to analyze the protein density in individual clusters. The local density is defined as the number of events present in a squared area of side 50nm centered on the event. The average density of events is defined as the arithmetic average of local density of the events composing the cluster. PAmCherry-PtsG clusters of population C exhibited significant lower average density values in presence of glucose (Fig. 9C), meaning that EII^glc^ complexes belonging to the same cluster localize further apart from each other when in presence of glucose, thereby occupying a larger membrane area.

## Discussion

Most bacteria transport various carbohydrates via a PTS system. The tremendous advantage of using a PTS coupled transport is that the substrate is phosphorylated in the process of import (hence the term group translocation), thereby effectively being removed from the chemical equilibrium of outside and inside concentrations. This allows the highly effective transport even under low external carbohydrate concentrations. Yet, a second, similarly important feature of PTS transport is that the complex PTS is composed of a sensory part and a regulatory part (4, 6). In enteric bacteria the EIIA^crr^ protein is a central component in the complex regulation cascade and a main part in the observed catabolite repression. EIIA^crr^ binds and inhibits the lactose permease as well as the GlpK protein in *E. coli*, while the phophsphorylated P~EIIA^crr^ activates the adenylate cycles (3). It is therefore not surprising that the EIIA component in *E. coli* is therefore a soluble protein that can dissociate from the membrane bound EIIBC complex. In line with this function subcellular localization of EIIA^Crr^ was shown to be dispersed in the cytoplasm (32). Also the general PTS components HPr is involved in regulation in *E. coli* and phosphorylates the transcriptional regulator BglG. For HPr a polar localization was described in *E. coli* that is alleviated when transport substrates are present (32). These data were in line with early suggestions that the PTS complex should act in multi-protein complexes (39), thereby improving its function (40). A major unsolved question in our understanding is still the unclear localization and assembly of the membrane embedded permease part of the PTS. Up to date there is only one report about the localization of an EII complex, the BglF in *E. coli* (32). However, BglF localization studies were performed with a plasmid borne expression system, potentially causing overexpression and therefore masking the possible clustering behavior of this PTS protein. While the general setup of proteins making a functional PTS is conserved in most bacteria, their genetic arrangement and regulatory role differs greatly among bacteria. In the high GC, gram positive *C. glutamicum*, four specific PTS were described (14). *C. glutamicum* differs greatly from organisms in which the PTS is well studied by the fact that it prefers utilization of several carbon sources simultaneously (9, 10). Hence, for most carbon sources *C. glutamicum* does not show diauxic growth behavior (except under conditions with ethanol or glutamate plus glucose). It is therefore not surprising that the permease subunits are fusion proteins composed of EIIABC and that they do not have a diffusible EIIA subunit. Our data using an N-terminal fluorescent fusion for the fructose specific EIIABC confirms clear membrane localization of the entire complex. It was generally assumed that the EII parts of the PTS are uniformly distributed in the cytoplasmic membrane, similar to other transport proteins, such as the Hxt hexose transporter in yeast (41), and the mentioned BglF (32). However, data on subcellular localization of transport proteins is rather scarce. This may in part be based on the relatively low copy number of many transport proteins that renders microscopy localization studies difficult. We have succeeded here in the subcellular localization of the EIIC permease parts of the fructose and glucose specific PTS. Both membrane integral transporters show a clustered membrane distribution. Importantly, both fusion constructs are fully functional based on their growth rates and the respective carbon sources consumption rates. Since the constructs replaced the native allele, we confidently assume also a wild type like copy number. Wide field microscopy not only revealed the heterogeneous, clustered localization of the two PTS components, but also showed that they hardly co-localize. Rather PtsG and PtsF seem to exclude each other. Interestingly, we could show that PtsF does co-localize with the succinate dehydrogenase, a protein of the TCA cycle and the respiratory chain. This finding indicates that in *C. glutamicum* proteins of the respiratory chain and transport proteins can co-occur in the same membrane region. The membrane economy model proposed by Zhuang *et al.* (42) suggests that bacteria regulate their membrane composition based on efficient usage of the limited membrane space and idealized a model in which a clear membrane separation of proteins involved in transport and respiration might occupy distinct membrane areas. For *C. glutamicum* we can exclude such a strict spatial distribution.

*C. glutamicum* PTS EII expression is known to be induced by the presence of the transported sugars (19), and regulation of the PTS gene expression is mainly controlled at the stage of transcription initiation or at transcription elongation (19). Our epifluorescence data support the induction of PtsF and PtsG in presence of the transported sugars by the increase in total fluorescence readings of cells under these conditions. Moreover, PtsF/G clusters increase in both size and foci area/cell area ratio, occupying more membrane space upon presence of glucose or fructose, while the overall number of large complexes, visible in widefield microscopy, decreases.

Importantly, patchy distribution of membrane proteins has been described for many signaling and scaffold proteins. Prime examples of membrane receptor clustering are the chemotactic receptors. A seminal publication describing the self-organization of the *E. coli* chemotaxis receptors using localization microscopy reveled that polar chemotaxis clusters mature by a stochastic assembly of smaller clusters and single receptor proteins (38). Other membrane proteins, such as flotillins (43, 44), the OXPHOS components Nuo, CydAB, CyoABCD, SdhABC (45, 46)(and own results, see Fig, 4E) have also been shown to localize in clusters of various size in the membrane. It is reasonable to assume that the clustering in these cases is based on stochastic self-assembly and that cluster formation is important for function. In contrast to scaffolding proteins such as flotillins (44, 47), it is not immediately apparent why a transport protein should cluster for an improved function. The PTS system however, is, as we described above, not only a transport system, but also a signalling device. Hence, clustering may be advantageous for signalling. In this context, it is interesting to note that a direct link between the PTS and the chemotaxis system has been described in *E. coli* (48, 49).

Here, we could show with single molecule localization microscopy, that the observed PTS cluster dynamically change their cluster density based on presence or absence of substrate. PAmCherry-PtsG PALM data showed that the presence of glucose in the medium induces expression and leads to cells exhibiting a higher number of larger clusters composed of 10<x<25 and 25<x events. These clusters showed a higher Cluster Max Span, but lower Average Density of Clusters values, meaning that the number of PtsG EII proteins around another decreases in presence of the transported sugar. At the same time, the distribution of number of events per cluster remained the same independently of the carbon source. These data strongly suggest a spatial rearrangement of PTS complexes, which might be a strategy to increase efficiency of membrane space utilization, or a form of regulation employed by cells (Fig. 10). Furthermore, a similarity between the function of chemotaxis receptors and the membrane bound EII complexes of the PTS is the crucial involvement of phosphorylation reactions. For the chemotaxis receptor it has been described that clustering improves phosphorylation reactions. Strikingly, it has also been described that different chemoreceptor densities lead to different kinase activity based on the local concentration of the receptors (50). In widefield microscopy these clusters are polar localized, but when cells treated with an excess of attractant, polar clusters disperse and proteins distribute over a large area of the membrane. It has been proposed that signal amplification under conditions of high attractant concentration is not needed (51). In a similar concept one might speculate that the PTS EII complexes in *C. glutamicum* increase their cluster density in absence or at very low concentration of the transport substrate. Under such conditions a positive cooperative effect might be useful. At higher concentrations of the transport substrate this cooperativity may not be advantageous and hence the PTS spreads over a larger membrane area. We observed a change in cluster size with PtsF and PtsG.

**Figure 10.**
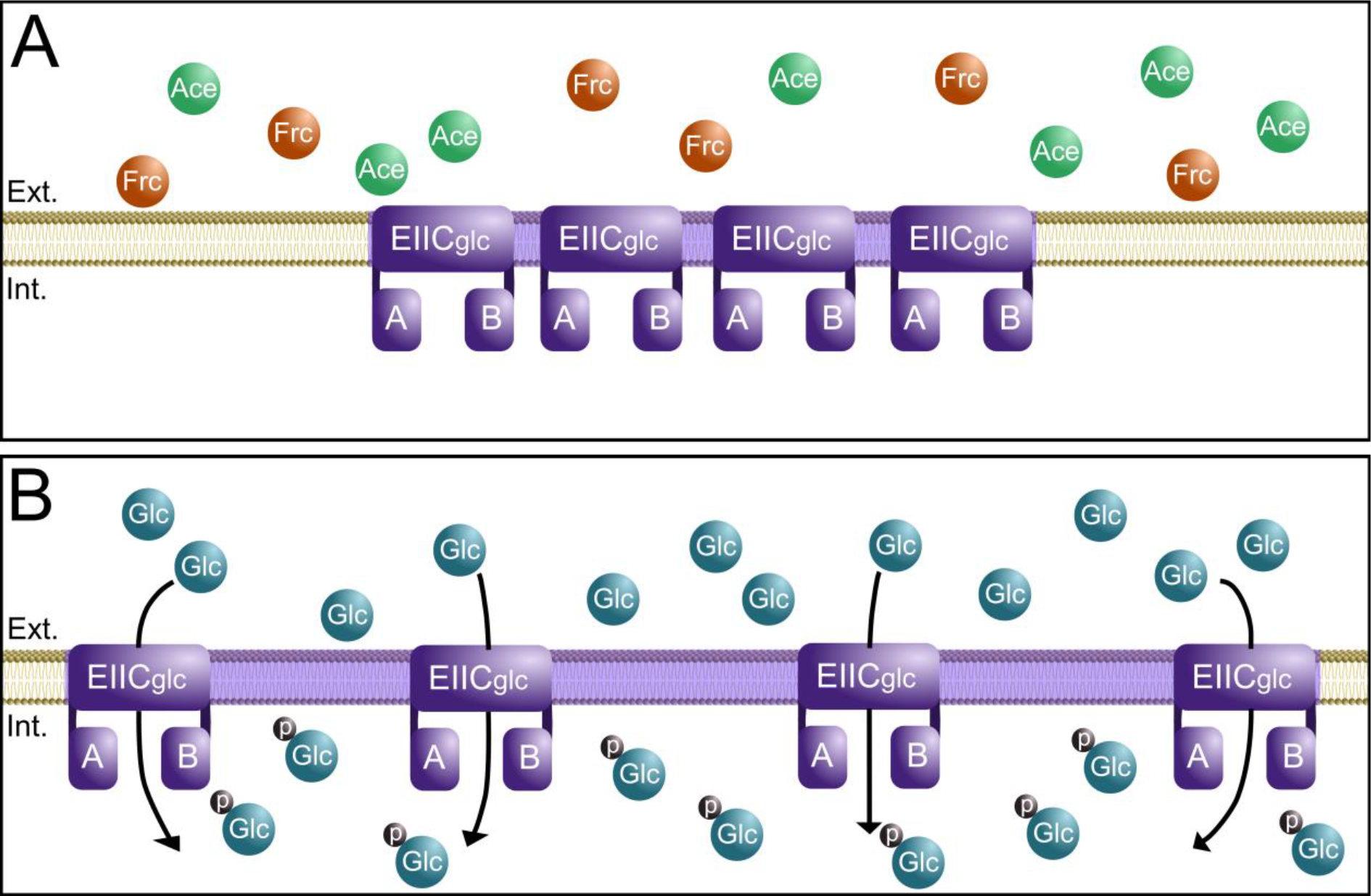
Membrane integral PtsG cluster undergo spatial rearrangement in presence of glucose. Schematic view of PtsG membrane occupancy in *C. glutamicum*. (A) In absence of the PTS sugar, in this case glucose, PtsG cluster are densely packed with proteins. These clusters occupy only a minimal membrane area. (B) In presence of the correct PTS substrate PtsG clusters rearrange, reducing the overall cluster density and occupying larger membrane areas.

However, statistically solid data were only achieved for PtsG due to the higher natural expression level. However, our widefield microscopy data further support this observation. This is the first time that a dynamic change in cluster density has been observed in cells and reveals an unexpected spatial regulation of the PTS. Several groups report data that suggest dimer formation of EII permeases (17, 52, 53). The nature of our experimental design in single molecule localization does not allow to distinguish between monomer and dimer in the membrane of *C. glutamicum* reliably.

Using artificial, induced L-form states, we could show that PTS cluster formation is not due to the rod-shape of the cells and the limited membrane surface. Although L-form cells show a grossly altered number of PTS clusters (likely owed to the higher expression as a result of the presence of multiple chromosomes), and also cluster size was larger, the same foci area/cell area ratio was retained, supporting the notion that membrane space is indeed a constraint that limits membrane occupancy by PTS proteins. However, clustering is an intrinsic function of the PTS EII.

Understanding the precise membrane coverage and spatio-temporal distribution of the PTS systems will help to model and scale up growth specific sugar uptake in industrial processes such as amino acid production in *C. glutamicum*. A kinetic model for glucose PTS in *E. coli* has been constructed (54, 55) and converted spatio-temporally to address the possible effects of diffusion in the PTS systems. Based on the model, it was concluded that the soluble E. *coli* EIIA^glc^ remains with up to 20% close to the membrane (55, 56). Thus in larger cells such as eukaryotic cells (or as shown here with L-forms) diffusion may disrupt efficient signaling due to diffusion limitation and subsequent gradient formation. However, these data were made under the assumption that membrane complexes as well as general PTS proteins are uniformly distributed either in the membrane (permeases) or cytosol (general PTS components). In the light of the data provided in this work, kinetic flux models might be adapted.

Currently, it remains unclear what triggers this remarkable change in cluster conformation. Future works need to discriminate whether substrate binding or the transport (e.g. phosphorylation) of the carbohydrates is required. We will address this question by placing mutations within the phosphotransfer reaction site of EIIAB or by using carbohydrate mimetics that bind EIIAB, but do not get transported. Furthermore, it remains to be tested, whether the observed dynamic cluster response to substrates is a peculiar finding in *C. glutamicum* or whether this could be a general mechanism found in other bacteria such as firmicutes or enteric bacteria.

## Material and methods

### Bacterial strains and cloning

Oligonucleotides, strains and plasmids used in this study are listed in Tables S1 and S2, respectively. DNA manipulations and *E. coli* DH5α transformations were carried out using standard cloning methods (57), and all constructed plasmids were verified by DNA sequencing.

### Media and growth conditions

For genetic manipulations, *E. coli* strain DH5α was grown at 37 °C in Lysogeny Broth (LB) medium supplemented, when appropriate, with kanamycin 25 μg/ml.

*C. glutamicum* cells were grown aerobically on a rotatory shaker (200 RPM) at 30°C in brain heart infusion (BHI) medium (Becton Dickinson) for maintenance, and in CGXII (58) supplemented with 2% of carbon source (glucose, fructose or acetate) for all microscopy and growth analysis. Acetate was used as a base carbon source, as it allows relatively good growth among non-PTS carbon sources in *C. glutamicum*. For all the experiments, strains were cultivated during the day in 5 ml of LB medium and then diluted in 10 ml of LB/CGXII medium and grown overnight. The following day the cultures were diluted to an optical density (OD) of 1.0 in 10 ml CGXII. For microscopy, samples were taken in early exponential phase, when the OD reached 2.

Bacterial L-forms were grown in osmoprotective medium composed of 2× MXM medium, pH 7 (40 mM MgCl_2_, 1 M xylose, and 40 mM maleic acid), mixed 1:1 with 2× CGXII supplemented with either glucose or fructose, depending on the sugar transported by the analyzed protein. As RES167 does not originally utilize xylose, this sugar was used for osmotic protection instead of sucrose to avoid possible influences in the studied PTS complexes.

### Carbon consumption assays

The quantification of glucose, fructose and acetate in the supernatants of cultures was performed by high-performance liquid chromatography (HPLC) using a Dionex UltiMate 3000 (Thermo Scientific) HPLC system equipped with an Aminex HPX-87H column (300 by 7.8 mm; Bio-Rad). Isocratic elution was performed with 6 mM H_2_SO_4_ at 80°C for 20 min at a flow rate of 0.6 ml/min. Fructose and glucose were detected with a PDA-100 Photodiode Array Detector (Dionex) at 190 nm, and acetate was detected with a PDA-100 Photodiode Array Detector (Dionex) at 206 nm. Quantification was performed using calibration curves obtained with external standards.

### Fluorescence microscopy

For standard fluorescence microscopy, cells were grown to early exponential phase as described before (59) and 1 μl of culture was spread on a 1% agarose gel pad. The setup used for fluorescent microscopy consisted of a Delta Vision Elite (GE Healthcare, Applied Precision) equipped with an Insight SSI^™^ illumination, an X4 laser module and a Cool Snap HQ2 CCD camera was used (100× oil PSF U-Plan S-Apo1.4 NA objective). Digital image analysis, was performed with Fiji (60). The corrected total fluorescence (CTF) was calculated according to following formula: CTF = Integrated Density - (Area of selected cell X Mean fluorescence of unspecific background readings) (61).

All statistical analysis and plotting were performed in RStudio and final image preparation was done in Adobe Photoshop CS2 (Adobe Systems Incorporated). All imaging experiments were performed several times with biological replicates, and PtsF/G foci analysis was performed in >200 cells.

### Photoactivated Localization Microscopy (PALM)

After growing to early exponential phase in CGXII supplemented with the desired carbon source as described before, cultures were fixed through incubation for 30 minutes at 30°C in formaldehyde in a final concentration of 1% v/v. Cells were then harvested by centrifugation and washed 3 times with PBS + 10 mM glycin, to be finally resuspended to 2 ODunits (OD.mL) in 200 μL TSEMS (50 mM Tris, pH 7.4, 50 mM NaCl, 10 mM EDTA, 0.5 M sucrose, 1x Protease Inhibitor Cocktail (Sigma), 13.2 ml H_2_O).

Imaging was performed using a Zeiss ELYRA P.1 equipped with the following laser lines: a HR diode 50 mW 405 nm laser and a HR DPSS 200 mW 561 nm laser and an Andor EM-CCD camera iXon DU 897 camera. Fluorescence was detected using a long pass 570nm filter (LP570), similar to a procedure described before (62).

An alpha Plan-Apochromat 100x/1,46 Oil DIC M27 objective was used for imaging. 100 nm TetraSpeck microsphere and the implemented drift correction tool were used to check for lateral drift and eventual drift correction. Calculation of the PALM image was performed via the 2D x/y Gaussian fit provided by the Zen2 software (Zeiss) with the following parameters: a peak mask size of 9 pixels (1 pixel = 100 nm) and a peak intensity to noise ratio of 6 (overlapping events were discarded). Z-axis was stabilized using the “Definite Focus” system.

The *C. glutamicum* strains *ptsF::pamCherry-ptsF* and *ptsG::PAmCherry-linker-ptsG* were imaged in the same way for each different condition: fructose, glucose and acetate. During the 10,000 frames that were collected during imaging, PAmCherry was activated using a linear gradient of the 405 nm laser ranging from 0.01 % to 10% (the 405 laser power was chosen in a way that minimized conversion of two separate PAmCherry molecules in close proximity at the same time). Converted PAmCherry was imaged using the 561 nm laser at 15 % (transfer mode) for 50 ms using an EMCCD gain of 200.

The localization events recognized by the Gaussian fit were filtered for photon number (70-350 photons) and point spread function (PSF) width at 1/e maximum (70-170 nm) in order to exclude the localization events originated by background (i.e.: dust particles) and/or the co-occurrence of multiple PAmCherry molecules.

### Western Blot and in-gel fluorescence

For western blot confirmation of the strains, cultures were grown in CGXII with 2% of the specific Pts sugars as carbon sources to an OD_600_ = 4, then harvested by centrifugation and resuspended in 1 mL of disruption buffer (100 mM NaCl, 100 mm KCl, DNAse, Protease inhibitor). Cells were subsequently disrupted by 10 cycles of 30 s at 6 m/s in FastPrep 24 (MP Biomedicals). The lysate was then centrifuged (18894 *g*, 15 min) and the supernatant containing the membrane fraction was mixed with loading dye and 15 μL was loaded on 0.1% SDS-containing 12% polyacrylamide gel, separated by electrophoresis and transferred to a PVDF membrane. Western blotting was carried out using standard methods: Blocking step was performed by incubation of the membrane in 10 ml TBS + 5% milk for 1h at RT, the primary antibody anti-mCherry was added in a 1:2000 dilution, followed by incubation for 1 hour. After washing the membrane 3 times with TBS, the secondary anti-rabbit antibody was added diluted 1:10000 to 10 ml TBS-T + 5% milk and incubated membrane for 1hr at RT. After another washing step, 5-bromo-4chloro-3-indolylphosphate and nitroblue tetrazolium solution (NBD/BCIP) was added and the membrane was incubated in the dark until the colors on the membrane were developed.

For in-gel fluorescence of strains containing mNeonGreen fusions, SDS gels cwere analyzed with a Typhoon Trio 9410 (Amersham Biosciences, GE) with 488nm laser excitation of mNeonGreen and fluorescence was detected using a 520nm filter.

## Acknowledgements

We thank Karin Schubert (LMU Munich) for the help with HPLC analysis.

## Author contributions

G.B.M., G.G. and M.B. designed the study

G.B.M. performed the genetic manipulations, strain characterization and widefield microscopy analysis

G.G. performed the PALM analysis

G.B.M., G.G. and M.B. analyzed the data and wrote the manuscript.

## Supplementary data

**Table S1.**
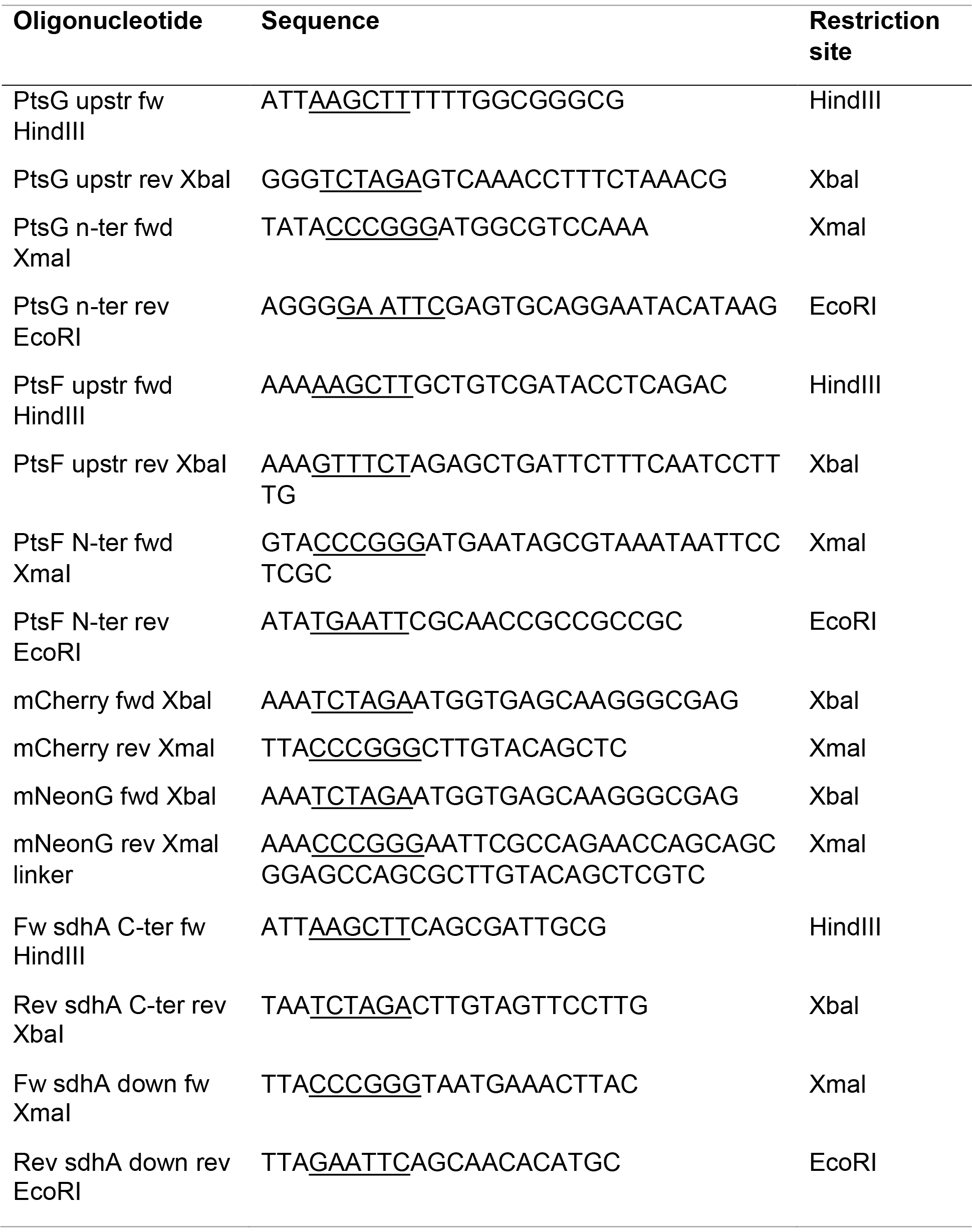
Oligonucleotides used in this study

**Table S2.** Strains and plasmids used in this study

| Strain or plasmid | Relevant characteristics | Source |
| --- | --- | --- |
| ***E. coli* strains** |  |  |
| DH5 $\alpha$ | F-Φ80lacZM15 (lacZYA-argF)U169 recA1 endA1 hsdR17(rk-,mk+) phoA supE44 thi-1 gyrA96 relA1 λ- | Invitrogen |
| ***C. glutamicum* strains** |  |  |
| RES 167 | Restriction-deficient derivative of strain ATCC 13032 | Tauch *et al.* (24) |
| CGM001 | RES167 derivative with *ptsF::mCherry-ptsF* | This work |
| CGM002 | RES167 derivative with *ptsG::mNeonGreen-ptsG* | This work |
| CGM003 | RES167 derivative with *ptsF::mCherry-ptsF* and *ptsG::mNeonGreen-ptsG* | This work |
| CGM004 | RES167 derivative with *ptsF::PAmCherry-ptsF* | This work |
| CGM005 | RES167 derivative with *ptsG::PAmCherry-Linker-ptsG* | This work |
| CGM006 | RES167 derivative with *sdhA::sdhA-mNeonGreen* |  |
| **Plasmids** |  |  |
| pK19mobsacB | Kan<sup>r</sup>; *E. coli*/*C. glutamicum* shuttle vector for construction of insertion and deletion mutants in *C. glutamicum* |  |
| pK19msB-mNeonGreen-ptsG |  | This work |
| pK19msB-mCherry-ptsF |  | This work |
| pK19msB-PAmCherry-ptsF |  | This work |
| pK19msB-PAmCherry-ptsG |  | This work |
| pK19msB-sdhA-mNeonGreen |  | This work |

**Figure S1.**
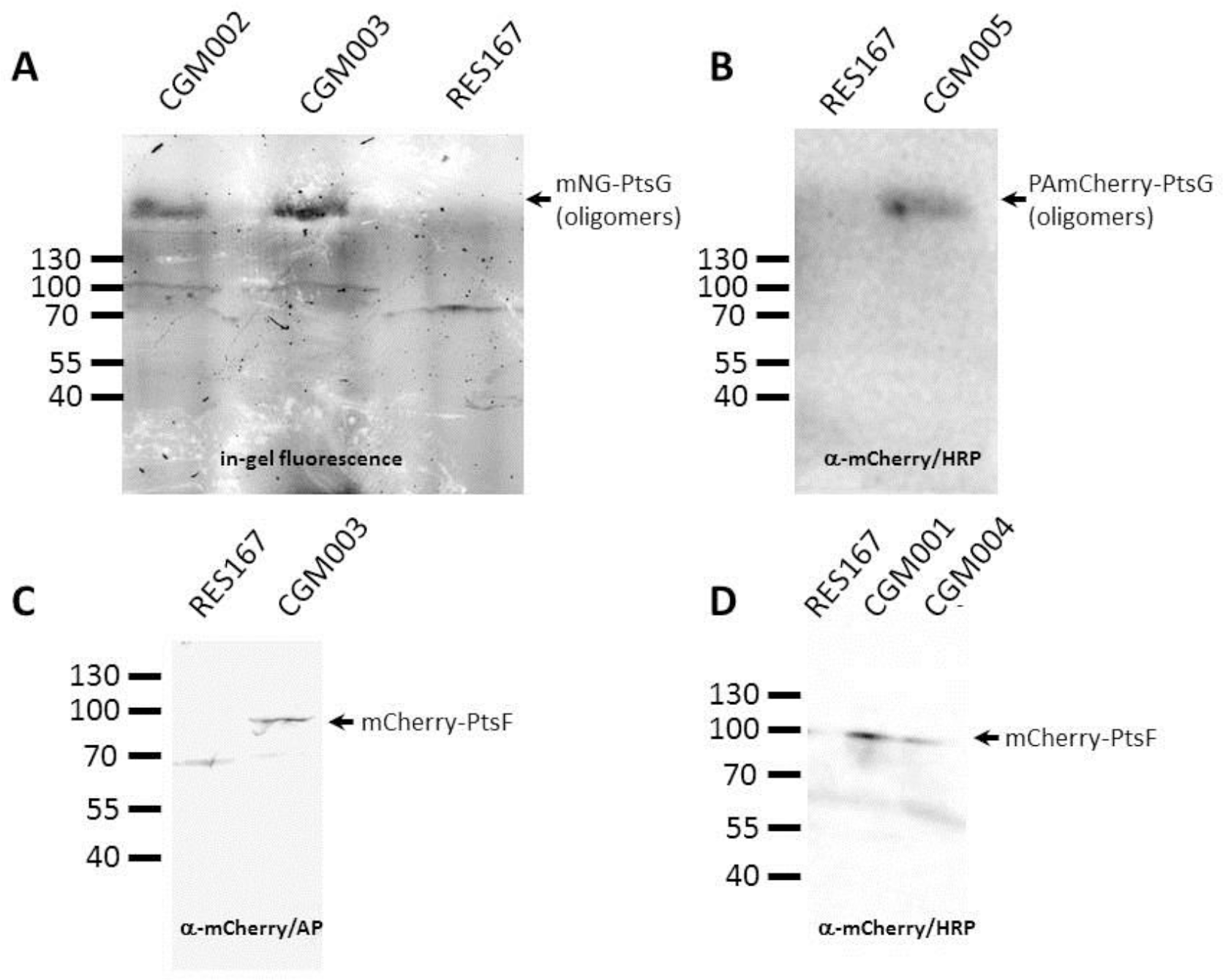
Control for full-length fusion proteins. In gel-fluorescence (A) and western blot analysis (B-D) with cell lysates of strains CGM001-006. (A) In gel fluorescence of cell lysates of CGM002 (mNeonGreen-PtsG) and CGM003 (mNeonGreen-PtsG in double labelled strain background) reveal the existence of oligomeric mNeonGreen-PtsG. (B) Western blot of cell lysate of CGM005 (PA-mCherry-PtsG) developed with α-mCherry antibodies and HRP-coupled secondary antibodies. PA-mCherry-PtsG is detected as oligomeric band with no apparent degradation products. (C) Western blot of cell lysate of CGM003 (mCherry-PtsF) developed with α-mCherry and AP-coupled secondary antibodies. (D) Western blot of cell lysate of CGM001 (mCherry-PtsF) and CGM004 (PA-mCherry-PtsF) developed with a-mCherry and HRP-coupled secondary antibodies. Note that little degradation is observed for all fusion constructs. PtsF = 70.51 kDa, PtsG = 72.57 kDa, mCherry = 28.8 kDa, mNeonGreen = 26.6 kDa.

**Figure S2.**
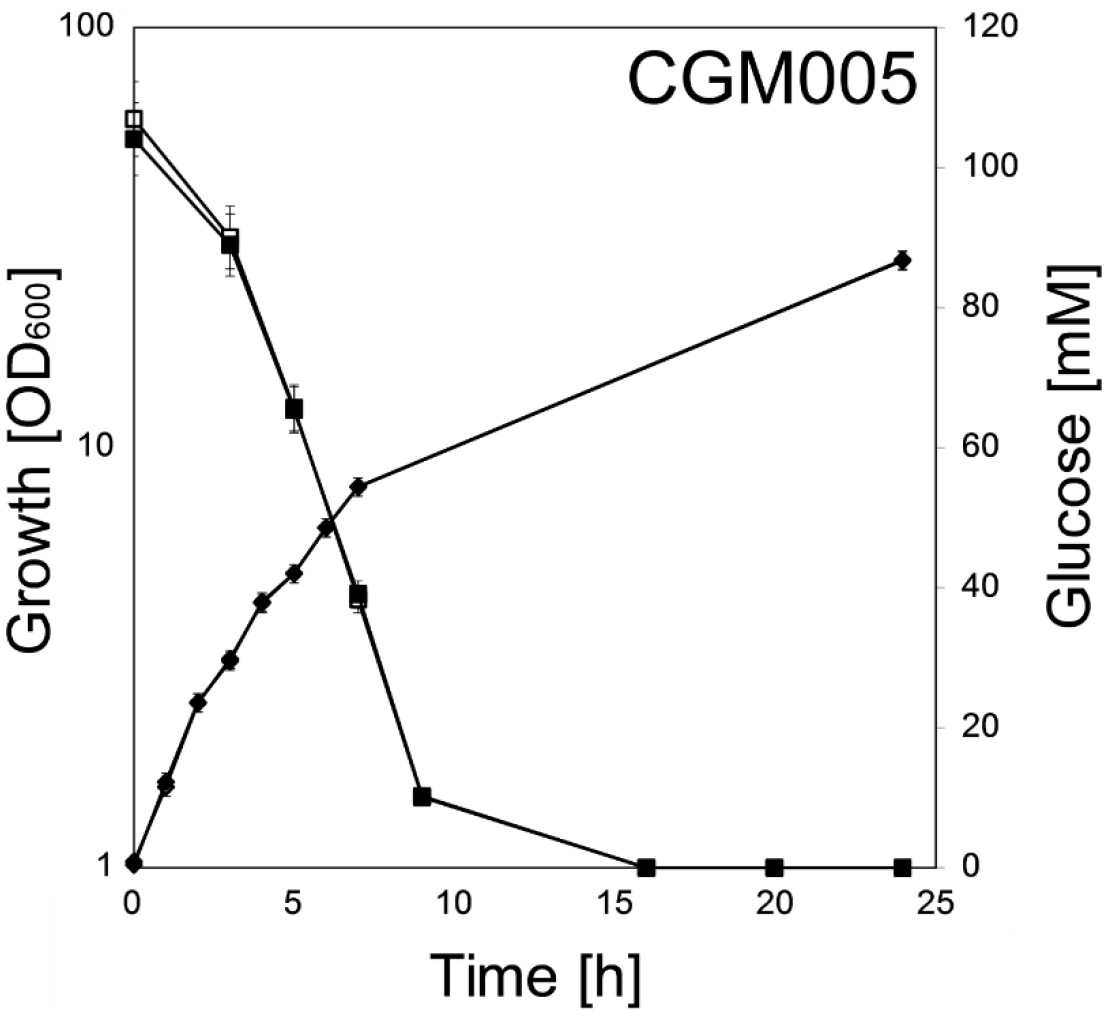
Growth and sugar consumption of *C. glutamicum* strain CGM005 (*PAmCherry-ptsG* filled symbols) versus wild type RES 167 (open symbols) on CGXII containing 100 mM glucose. Glucose consumption (squares) and growth (diamonds) are indicated. Each point represents biological triplicates and standard deviation is indicated.

**Figure S3.**
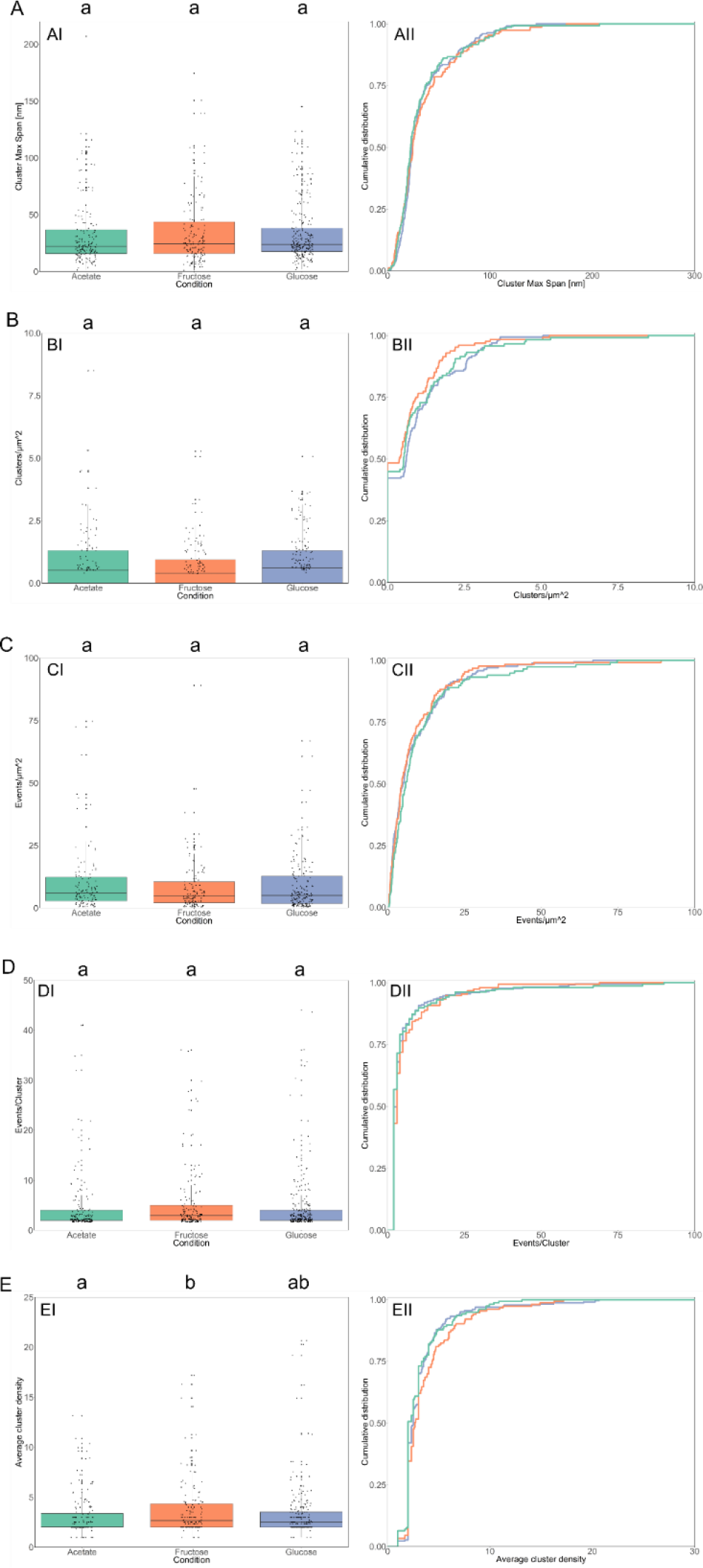
Statistical analysis of single molecule resolution PALM data of *C. glutamicum* strain *ptsF::PAmCherry-ptsF* under different growth conditions. (A) PAmCherry-PtsF clusters Maximum Span, (B) PAmCherry-PtsF clusters per μm^2^, (C) number of events per μm^2^, (D) number of PAmCherry-PtsF events in clusters, (E) average density of clusters composed of >2 molecules: the arithmetic average of Local Density of the events composing a cluster. Cells were grown in CGXII supplemented with 2% of indicated carbon sources.

**Figure S4.**
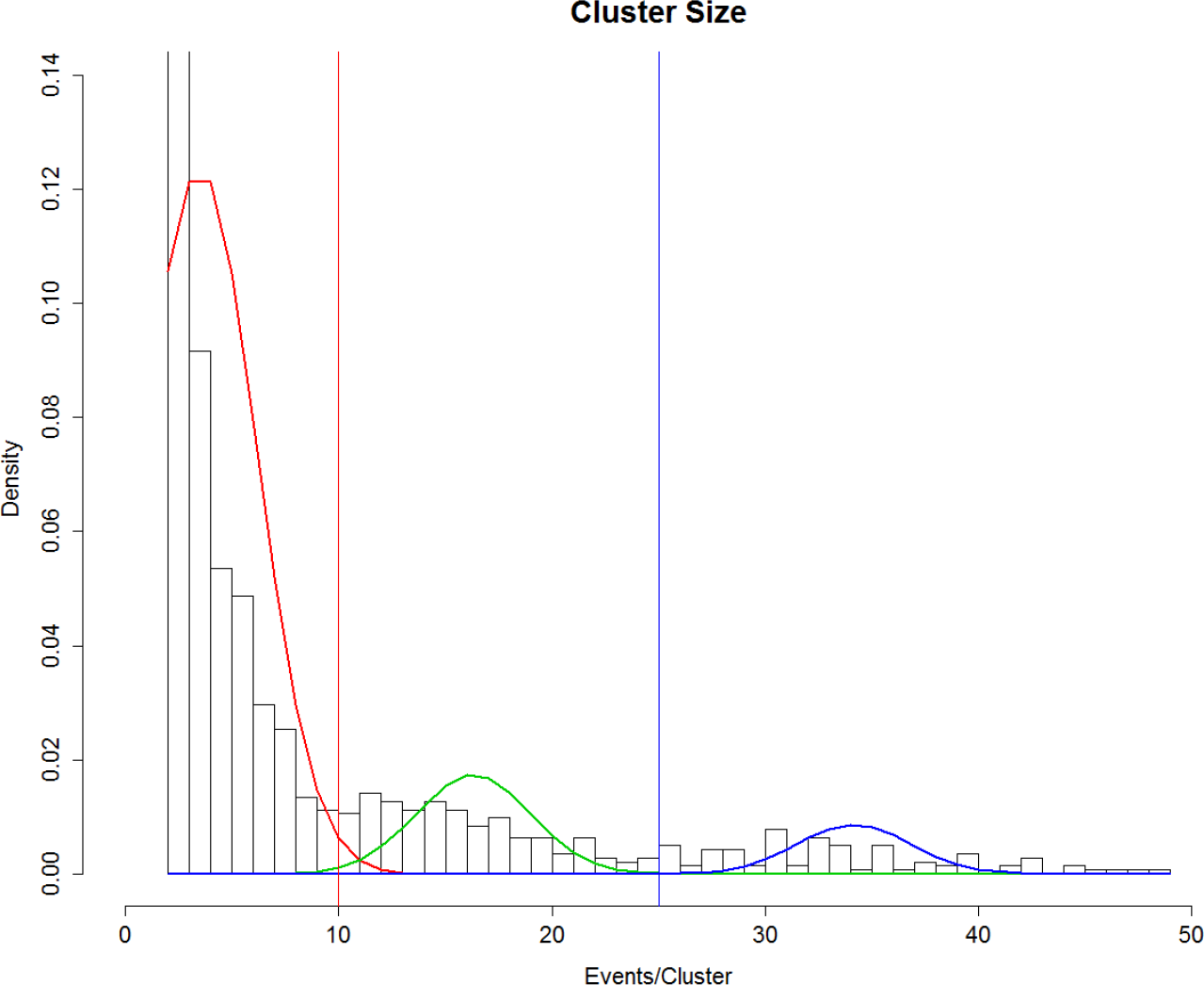
PtsG-PAmCherry clusters size distribution. Three different populations were identified by univariate normal mixture analysis: clusters composed of 2-9, 10-24, >25 events. Lines of different colors represent Gaussian fits for each different population.

